# The NF-κB pathway controls H3K9me3 levels at intronic LINE-1 elements and hematopoietic stem cell gene expression in cis

**DOI:** 10.1101/2021.06.08.447574

**Authors:** Yanis Pelinski, Donia Hidaoui, Anne Stolz, François Hermetet, Rabie Chelbi, M’boyba Khadija Diop, Amir M. Chioukh, Françoise Porteu, Emilie Elvira-Matelot

**Affiliations:** Inserm UMR1287, Université Paris-Saclay, Gustave Roussy, 94805, Villejuif, France; Bioinformatics Platform UMS AMMICa INSERM US23/CNRS 3655, Université Paris-Saclay, Gustave Roussy, 94805, Villejuif, France

**Author notes:** to whom the correspondence should be sent: Françoise Porteu, Emilie Elvira-Matelot, INSERM U1287, Cancer Campus Gustave Roussy, PR1, 114 rue Édouard-Vaillant, 94805 Villejuif Cedex-France. Equal contribution (Donia Hidaoui, Anne Stolz, Françoise Porteu, Emilie Elvira-Matelot).

**Keywords:** hematopoietic stem cells, irradiation, heterochromatin, retroelements, TNF-α, NF-κB

## Abstract

Ionizing radiations (IR) alter hematopoietic stem cell (HSC) function on the long-term, but the mechanisms underlying these effects are still poorly understood. We recently showed that IR induces the derepression of L1Md, the mouse young subfamilies of LINE-1/L1 retroelements. L1 contribute to gene regulatory networks. However, how L1Md are derepressed and impact HSC gene expression are not known. Here we show that IR triggers genome-wide H3K9me3 decrease that occurs mainly at L1Md. Loss of H3K9me3 at intronic L1Md harboring NF-κB binding sites motifs but not at promoters is associated with the repression of HSC specific genes. This is correlated with reduced NFKB1 repressor expression. TNF-α-treatment before IR rescued all these effects and prevented IR-induced HSC loss of function *in vivo.* This TNF-α/NF-κB/H3K9me3/L1Md axis might be important to maintain of HSCs while allowing expression of immune genes during myeloid regeneration or damage-induced bone marrow ablation.

## INTRODUCTION

Exposure to ionizing radiations (IR), in the context of medical use such as radiotherapies, is an independent risk factor for many disorders characteristic of an accelerated aging. The hematopoietic tissue is particularly sensitive to IR.

HSCs self-renew and give rise to all blood cell lineages. Maintenance of their integrity is crucial throughout life. We and others have shown that total body irradiation (TBI) in mice leads to long-term defects in hematopoiesis due to loss of HSC reserves and functions (Fleenor et al., 2015; de Laval et al., 2013; Mohrin et al., 2010). In HSCs, IR induces DNA damage accumulation, loss of self-renewal, and a biased differentiation towards the myeloid lineage leading to increased myeloid cell counts and decline of the adaptive immune response. These changes are likely contributing to many IR-induced premature aging disorders and to the high risk of developing myeloid leukemia. Understanding the molecular mechanisms leading to HSC loss of function upon IR is necessary to modulate its adverse effects. It may also help identifying the first events leading to hematologic malignancies.

IR induces DNA double strand breaks (DSBs). DNA damage is thought to be one of the main driving forces of aging. However, delaying the effects of age in mice by decreasing the levels of DNA damage has never been achieved, and a direct link between DSB formation and physiological aging is still lacking (White and Vijg, 2016). Although IR has been shown to induce chromosomal abnormalities in progenitors (de Laval et al., 2013, 2014; Mohrin et al., 2010), in fact HSCs are quite resilient towards accumulating DNA mutations in response to DNA damage (Moehrle et al., 2015).

In addition to DSBs, IR has been shown to induce changes in chromatin state, mainly at the level of DNA methylation, in different tissues and cell lines (Miousse et al., 2017b). Epigenetic alterations have been observed in aged HSCs in the absence of mutations in epigenetic factors (Djeghloul et al., 2016; Sun et al., 2014). Reorganization of heterochromatin is among the most commonly-reported changes in aging and senescence supporting its essential role in maintaining proper cellular function (Tsurumi and Li, 2012). Maintenance of HSC identity is dependent on the heterochromatin mark H3K9me3 (Koide et al., 2016). Decreased H3K9me3 in HSCs due to loss of *Suv39h2* and/or *Suv39h1* methyltransferases occurs with age and is associated with the loss of B cell differentiation (Djeghloul et al., 2016) and hematopoietic changes archetypal of aging (Keenan et al., 2020). However, whether the long-term effects of IR on HSCs is linked to IR-induced changes in heterochromatin remains to be addressed.

Heterochromatin plays also a major role in the maintenance of the genome stability by repressing transposable elements (TEs), including DNA transposons, and retrotransposable elements (RTEs), further classified as long terminal repeat (LTR) sequences which characterize endogenous retroviruses (ERV), and non-LTR elements such as long or short interspersed elements (LINE-1/L1; SINE). Propagation of RTE in the genome has been recognized as a great source of genomic instability. Even without propagating, RTE have also been recently recognized as major contributors of gene regulatory networks (Chuong et al., 2017). Indeed, they qualitatively and quantitatively control gene expression, providing alternative enhancers, promoters, splicing or polyadenylation signals, and also serving as cis-regulatory elements in a cell specific fashion. Basal L1 expression in early mouse embryo is necessary for its proper development (Jachowicz et al., 2017). RTE are involved in T-cell differentiation by regulating genes involved in immune processes (Adoue et al., 2019). Abnormal RTE expression has been observed in cancers, including AML, and may be involved in the pathogenesis through the alteration of host gene expression (Chuong et al., 2017) and the expression of oncogenes (Deniz et al., 2020; Jang et al., 2019).

We recently showed that evolutionary recent mouse L1 (L1Md) are highly expressed in HSCs and that their expression is further increased and maintained at high levels up to 1 month after TBI (Barbieri et al., 2018). TE expression is also increased in HSC after chemotherapies (Clapes et al., 2021) and the decreased H3K9me3 in aged HSCs is associated with increased L1Md expression (Djeghloul et al., 2016). However, the impact of L1Md derepression on the HSC transcriptome remains to be addressed. In addition the mechanisms and signaling pathways by which IR specifically triggers L1Md expression in HSCs is currently unknown. We and others previously showed that the NF-κB signaling pathway is required to prevent IR-induced HSC injury (Hu et al., 2021; de Laval et al., 2014). In addition, tumor necrosis factor α (TNF-α)-induced NF-κB supports HSC survival during inflammation and chemotherapeutic stress induced by 5-fluorouracil (Yamashita and Passegué, 2019). On the other hand, basal activation of NF-κB is required for HSC homeostasis and self-renewal potential and the expression of key HSC maintenance genes are severely impaired in mice deficient for NF-κB pathway factors (Fang et al., 2018; Hu et al., 2021; Stein and Baldwin, 2013).

We show here that IR induces a major loss of H3K9me3 in HSCs, which mainly affects L1Md subfamilies, and more specifically intronic L1Md displaying NF-κB binding sites. By controlling the levels of H3K9me3 at selected intronic L1Md located in genes belonging to the HSC signature, this pathway plays a crucial role to preserve HSC specific gene expression and HSC function during IR stress.

## RESULTS

### H3K9me3 is mainly enriched at recent L1Md subfamilies in HSCs

To assess the effect of IR on H3K9me3 in HSCs and more particularly at TE sequences, we performed H3K9me3 ChIP-seq experiments in HSCs (Lin^−^Sca^+^-Kit^+^-CD34^−^Flk2^−^) sorted from mice one month after TBI (2Gy) or not, as previously described (Barbieri et al., 2018) (**Figure 1A-B**). Deep characterization of H3K9me3 enrichment in HSCs, notably at TE sequences, has never been performed. We thus first characterized H3K9me3 genomic coverage in HSCs at steady state.

**Figure 1.**
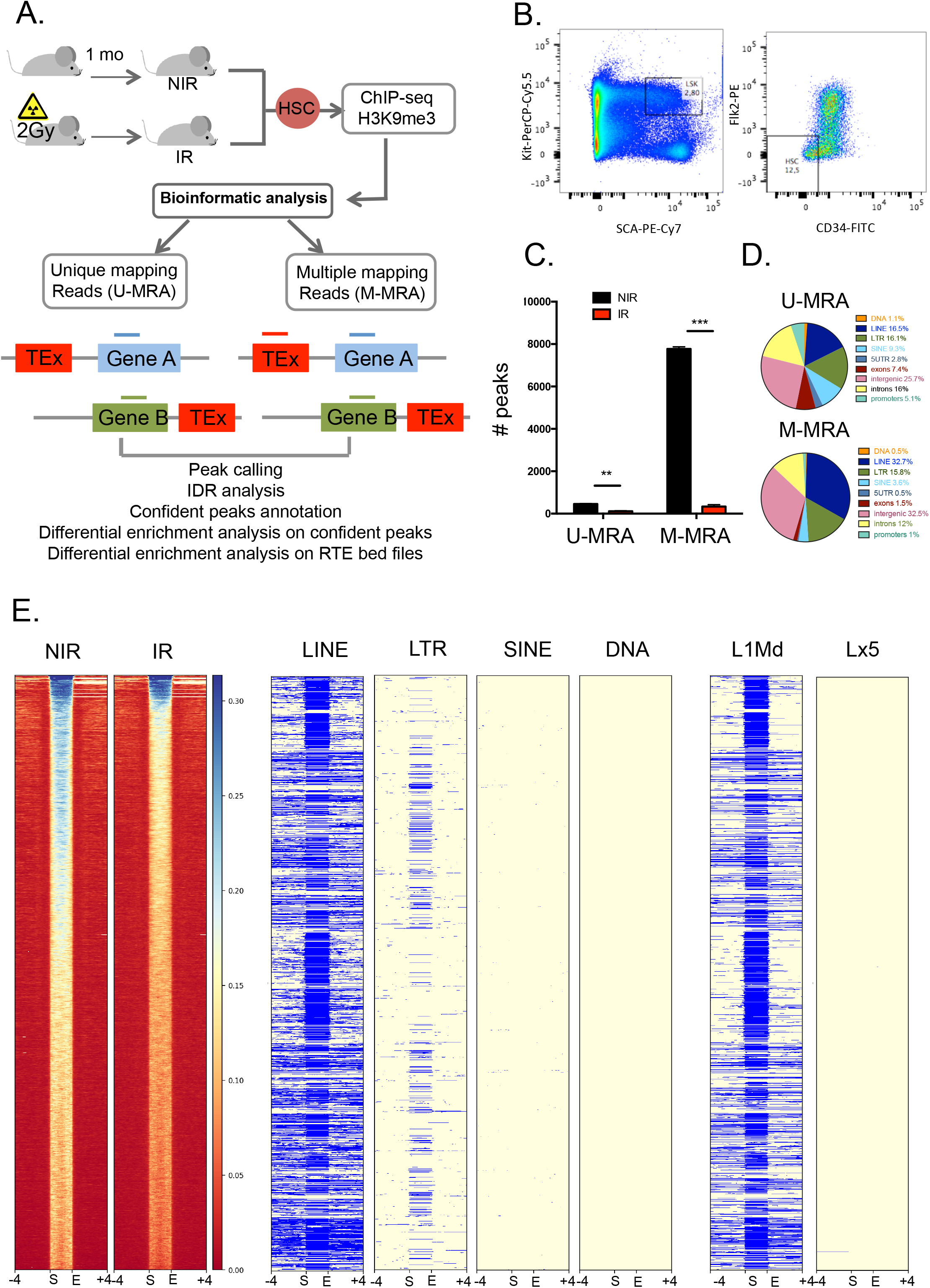
H3K9me3 is mainly enriched at recent L1Md subfamilies in HSCs. **(A)** Experimental and bioinformatic analysis design for H3K9me3 ChIP-seq. Mice were subjected to 2Gy total body irradiation (IR) or left untreated (NIR). Analysis was done on both Unique and Multiple mapping reads (U- and M-MRA respectively). (B) Gating strategy for HSCs (LSK CD34-Flk2-) sorting. (**C)** Number of peaks with IDR score <0.05 in U- and M-MRA and in NIR and IR conditions. (**D)** Repartition of confident peak annotations in each genomic feature using annotateR in NIR conditions for U- and M-MRA. *** p<0.001; ** p<0.01, t-test **(E) left columns** : Heatmap of H3K9me3 enrichment at M-MRA confident peaks in NIR and IR conditions. Each row represents one-scaled H3K9me3 peak with +/− 4kb flanking regions. The color scale represents H3K9me3 enrichment over the input (H3K9me3 concentration) with the red corresponding to lower enrichment, and the blue to stronger enrichment. **Center and right columns:** Genomic coverage of the H3K9me3 peaks showed in the heatmap at different TE families and subfamilies (LINE, LTR, SINE, DNA, L1Md, Lx5) are also represented. Blue, TE is present; yellow, TE is absent in the peak (row).

Analysis of genomic repeats is still a bioinformatics challenge. The assignment of reads complementary to sequences that are repeated in the genome and that present low sequence variation is largely compromised. These multiple mapping reads are often discarded in ChIP-seq studies. However, when using only unique mapping reads, one can induce a bias of representation of the TE subfamilies towards the oldest ones as the youngest subfamilies, such as L1Md, present a very low mappability due to quasi-identical copies (Teissandier et al., 2019). Therefore, while young families of TEs are the most epigenetically regulated in the genome (Barau et al., 2016; Pezic et al., 2014), they are severely underestimated in unique read analysis.

To maximize the output information on these sequences, we considered all reads that mapped to the genome without mismatches and randomly assigned them at one of their best possible position in the genome (**Figure 1A**). We combined both unique and multiple mapping reads analyses (U-MRA and M-MRA, respectively) to finally get a compromise between precise assignment of the unique reads and global information at youngest TE subfamily level.

Quality control of the resulting reads indicated high genomic coverage in both U- and M-MRA and in NIR and IR conditions **(Table S1)**. Peak calling followed by reproducibility measurement between replicates (Irreproducible discovery rate=IDR) identified 456 peaks on average in the NIR conditions in the U-MRA. As expected, with 7769 peaks, M-MRA gave a substantial increased number of peaks (**Figure 1C; Table S1)**. Our peak calling retrieved the SUV39h1 and 2-dependent H3K9me3 peaks that were previously described at young TEs such as IAP ERVs (**Figure S1A**) using this strategy in mouse embryonic stem cells (mESCs) (Bulut-Karslioglu et al., 2014). Of note, these elements were not covered by U-MRA.

After annotation of the peaks using annotateR, both U-MRA and M-MRA showed that the majority of the H3K9me3-enriched peaks occurs at TEs (41.9% and 52% respectively) (**Figure 1D**). Interestingly, M-MRA shows a gain in LINE representation, with 32.7 % of the total H3K9me3 enriched peaks *vs* 16.5% in U-MRA. By contrast, M-MRA shows reduced SINE and DNA representation while LTR representation was not affected. This suggests that H3K9me3 enrichment occurs mainly at LINEs in HSCs. Heatmap representation of the average concentration (*ie*: H3K9me3 enrichment normalized to input) of H3K9me3 at peaks retrieved from M-MRA, followed by RTE genomic coverage, further confirmed that peaks enriched in H3K9me3 at basal are mostly covering LINEs compared to LTR, SINEs or DNA, and more especially the young subfamilies of LINEs (L1Md) compared to older ones (Lx5) (**Figure 1E**).

Altogether, these data demonstrate that H3K9me3 is mainly enriched at repetitive sequences, and more particularly at young L1Md elements in HSCs.

### Irradiation induces a loss of H3K9me3 that mainly affects the recent L1Md subfamilies

We next investigated changes in H3K9me3 that occur upon IR. We first compared the number of H3K9me3 peaks identified in IR *vs* NIR conditions. Only 115 peaks in average were found in IR in the U-MRA, compared to 456 peaks in NIR condition, and 283 compared to 7769 in the M-MRA (**Figure 1C**). Heatmap representation further showed a loss of H3K9me3 enrichment at peaks retrieved from M-MRA upon IR (**Figure 1E**).

Differential H3K9me3 enrichment analysis performed at H3K9me3 peaks identified in both IR and NIR conditions further revealed a strong decrease in H3K9me3 enrichment in IR (**Figure 2A**). We found 13 and 253 peaks showing significant (*p<0.05*) differential H3K9me3 enrichment upon IR in U- and M-MRA, respectively. All of them in U-MRA and 252/253 peaks in M-MRA showed decreased H3K9me3 upon IR (**Figure 2A**). These data reveal a major loss of H3K9me3 genomic coverage upon IR, regarding both the number of peaks and the concentration of H3K9me3 at conserved peaks.

**Figure 2:**
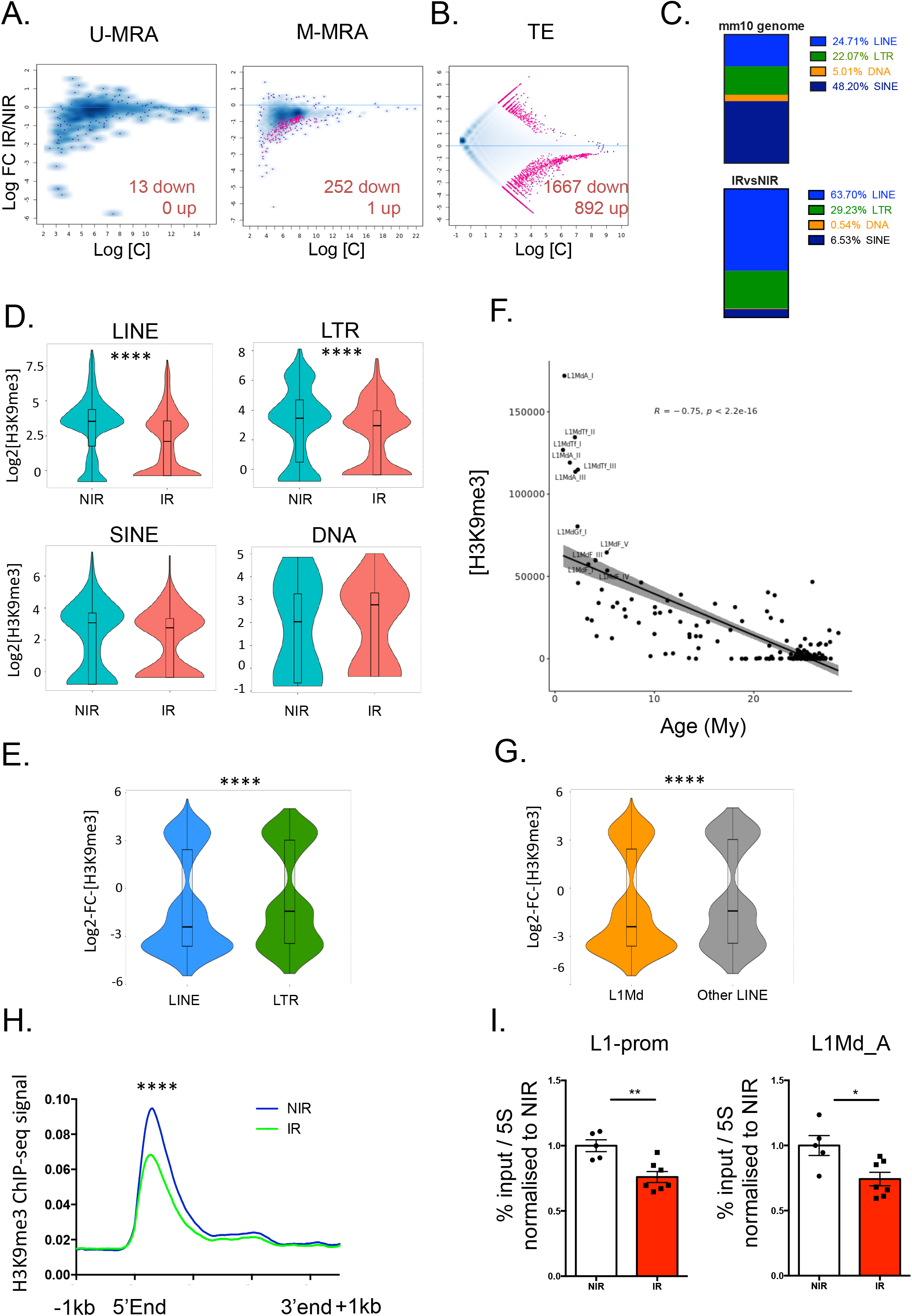
Irradiation induces a loss of H3K9me3 that mainly affects the recent L1Md subfamilies. **(A-B)** MA-plots showing non-significant (blue dots) and significant (*p<0.05-*pink dots) differential H3K9me3 enrichment at **(A)** confident peaks between NIR and IR conditions analyzed both in U and M-MRA and **(B)** TE loci. The number of peaks showing a significant decreased (down) or increased (up) in H3K9me3 enrichment upon IR is indicated in the plot. (**C)** Distribution of the % of each family of TE among the total TE loci in the mouse mm10 genome (up) and among the significantly differentially enriched TE (bottom) retrieved in (B). (D) Violin plot representing the distribution of H3K9me3 concentration at each locus retrieved in (B) for LINE, LTR, SINE and DNA families of TEs in NIR and IR conditions. **(E)** Violin plot representing the distribution of the log2 fold change in H3K9me3 concentration at each locus retrieved in (B) for LINE and LTR. (**F)** Correlation plot representing H3K9me3 concentration quantified at all LINE elements in M-MRA *vs* their age in million years (My). R, Pearson correlation coefficient; p, pvalue. **(G)** Violin plot representing the distribution of the log2 fold change in H3K9me3 concentration at each locus retrieved in (B) for L1Md or other LINE. **(H)** Plot profile of H3K9me3 enrichment along the L1Md sequences (>5kb) +/− 1kb flanking regions in IR (green) *vs* NIR (blue) conditions. (**I)** H3K9me3 enrichment at L1Md promoters analyzed by ChIP-qPCR normalized to H3K9me3 enrichment at repetitive 5S rRNA. n=3 independent experiments. Each dot represents pools of 3 (NIR) or 4 (IR) mice. Results are expressed as fold change from the mean value of the NIR condition and represented as means +/− SEM (*p<0,05; **p<0.01 t-test). **(D), (E), (G):** **** p<0.0001 t-test; **(H)** ****p<0.0001 wilcoxon test

We performed the same analysis at TE genomic loci instead of peaks and found 2559 loci with significant H3K9me3 differential enrichment, 1667 and 892 loci showing decreased or increased H3K9me3, respectively upon IR (**Figure 2B).** Annotation of these loci show that they are enriched in LINEs (66.7%) and LTR (29.23%) compared to the distribution of these TEs in the mouse genome (24.71 and 22.07 % respectively) (**Figure 2C**). The distribution of the H3K9me3 concentration showed a significant (*p<0.0001*) decreased in H3K9me3 at both LINEs and LTR upon IR (**Figure 2D).** Comparing the distribution of the log2 fold-change between IR and NIR conditions in H3K9me3 concentration showed that this decrease is significantly (*p<0.0001)* more pronounced at LINEs than at LTRs (**Figure 2E**).

Since young TE subfamilies, notably L1Md, were previously reported to be the most epigenetically regulated in the genome compared to old LINEs in ESCs and testis (Barau et al., 2016; Pezic et al., 2014), we further dissected H3K9me3 enrichment at TEs depending on their age, as calculated in (Sookdeo et al., 2013). We observed a significant negative correlation between the age of the LINE and H3K9me3 enrichment, with the youngest, typically L1Md_A and L1Md_Tf, showing the highest enrichment (**Figure 2F**), as previously observed for DNA methylation (Miousse et al., 2017a). IR induced a significantly (*p<0.0001*) more pronounced loss of H3K9me3 at young L1Md compared to other LINEs (**Figure 2G**).

Altogether these data reveal that one month after IR, HSCs display a major loss of H3K9me3, which mainly occurs at L1Md, the subfamilies of TEs the most enriched in H3K9me3 at steady state. This prompted us to further focus our analysis on L1Md.

Plot profile of H3K9me3 enrichment along L1Md sequences showed asymmetric distribution of H3K9me3 along the L1 body, with the highest enrichment and the highest loss upon IR at the 5’ end of these elements (**Figure S1B**). L1 retrotransposition is often abortive and results in the insertion of copies truncated in their 5’ end. Thus, a majority of L1 does not harbor the 5’ end. Plot profile centered on the longest L1Md (≥5kb) enriched in L1 harboring promoter sequences further confirms that IR mainly decreases H3K9me3 levels at this location (**Figure 2H**). Finally, we confirmed the global decrease of H3K9me3 at L1Md promoters, and more particularly at L1Md_A, using ChIP-qPCR experiments with primers recognizing either all L1Md or specifically L1Md_A promoter (**Figure 2I**), as described previously (Barbieri et al., 2018).

### Irradiation induces a strong deregulation of the HSC transcriptome

In order to unravel the consequences of IR-induced H3K9me3 loss on the HSC transcriptome, we performed RNA-seq of HSCs one month after TBI. Comparison of IR *vs* NIR revealed 1067 differentially expressed genes (DEGs) (*p<0.05*), with 602 (56.4%) genes downregulated and 465 (43%) genes upregulated upon IR (**Figure 3A and B**; **Table S2**). Differences in gene expression are very strong, as almost 80% of the DEGs between IR and NIR present a fold change above 10 and more than 50% present a fold change above 50 (**Figure 3B**). Gene Set enrichment analysis (GSEA) on Hallmark gene sets indicated significant enrichments in DNA repair, G2/M checkpoint and oxidative phosphorylation pathways, as expected upon IR (**Figure 3C**), whereas the main pathways lost in IR are related to cell signaling (**Figure 3D**). Among these, the most significant decrease concerns genes regulated by NF-κB in response to TNF-α (**Figures 3D and 3E**). A recent report showed that TNF-α induces a specific prosurvival gene signature in HSCs (Yamashita and Passegué, 2019). Interrogating this gene signature, composed of 62 genes representing both core regulators of the NF-κB pathway and TNFα-induced HSC-specific survival genes, we observed its significant loss upon IR (**Figure 3F**). IR also induced the loss of the different HSC TNF-α signatures taken individually: the two *in vitro* signatures obtained 3h and 12h after TNFα treatment and the *in vivo* signature obtained 3h after TNFα treatment in mice (**Figure S2A**). By contrast, IR had no effect on the GMP TNF-α gene signature (**Figure 3G**) (Yamashita and Passegué, 2019).

**Figure 3.**
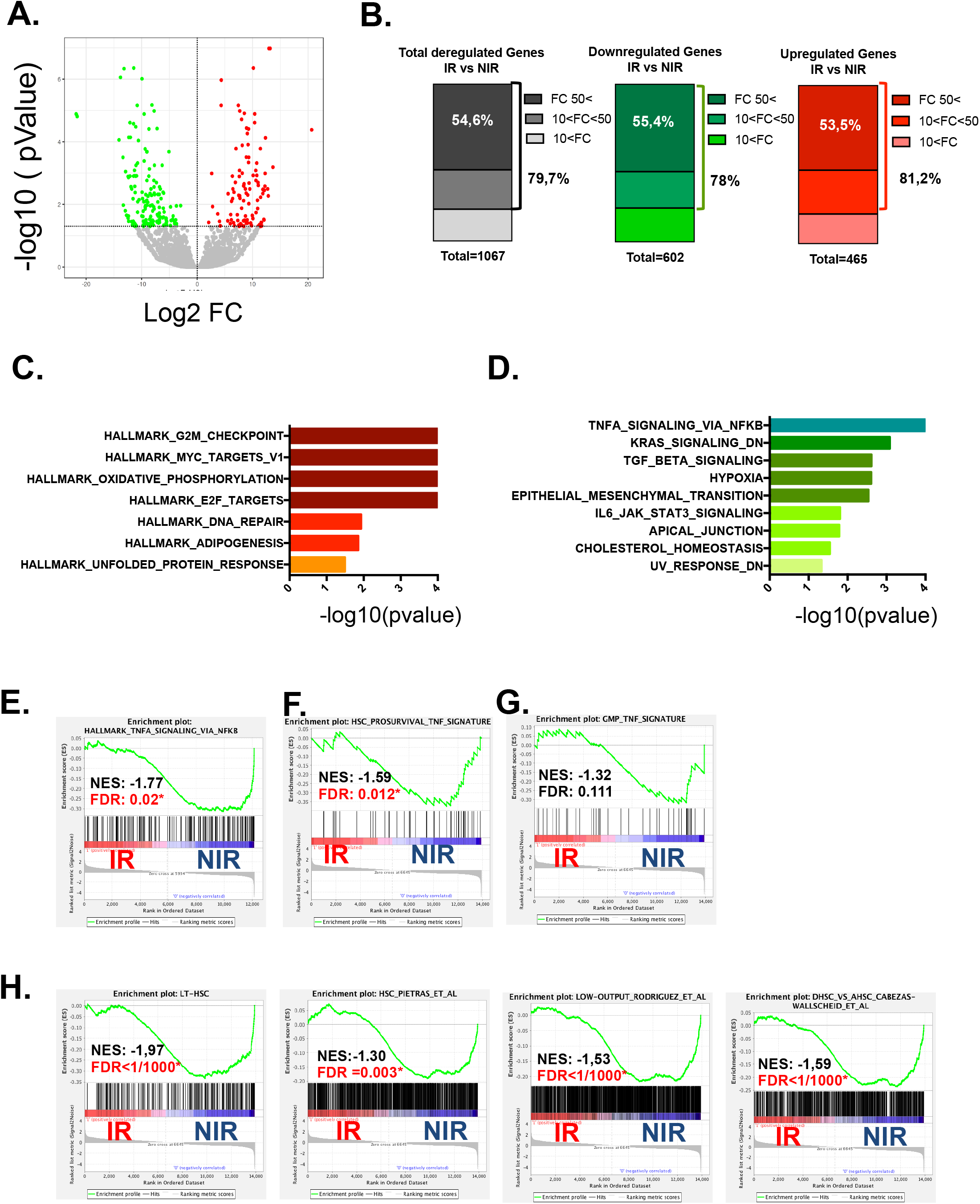
Irradiation induces a strong deregulation of the HSC transcriptome. **(A)** Volcano plot of the DEG between IR and NIR conditions. The horizontal axis represents the log2 fold change (FC) and the vertical axis the –log10(pValue). Significantly (*p<0.05*) upregulated (red) and downregulated (green) genes are shown. (**B)** Repartition of total (*p<0.05)* DEG (grey); downregulated (green) or upregulated (red) genes in IR *vs* NIR according to their fold change. (**C-H)** GSEA analysis using Gene Sets. Significant (*p<0.05*) gene sets gained (**C**) or lost (**D**) in the IR condition compared to NIR. log10(pValue) is set to 4 when p<0.001. Enrichment plots for TNF-α_signaling_via_NFKB hallmark (**E),** HSC prosurvival TNF-α (**F)** and GMP TNF-α (**G)** gene signatures. (**H)** Enrichment plots for LT-HSC; Low-output and dormant *vs* activated HSC signatures.

Given the importance of the NF-κB pathway at maintaining HSC survival and self-renewal or during chemotherapeutic and IR stress (Hu et al., 2021; de Laval et al., 2014; Yamashita and Passegué, 2019), we next interrogated different published HSC signatures that are enriched in low-output/self-renewing and functional long-term regeneration HSCs compared to differentiating HSCs, or in dormant *vs* activated HSCs (Cabezas-Wallscheid et al., 2017; Chambers et al., 2007; Pietras et al., 2015; Rodriguez-Fraticelli et al., 2020). We found a significant loss of all these signatures upon IR (**Figure 3H and S2B)**. Likewise, the megakaryocyte (MK)-Biased output HSC signature (Rodriguez-Fraticelli et al., 2020), representing platelet-primed HSCs that were also previously described to be at the apex of the HSC hierarchy (Sanjuan-Pla et al., 2013), was also enriched in NIR *vs* IR HSCs **(Figure S2B).** Conversely, the high-output and multilineage signatures (Rodriguez-Fraticelli et al., 2020), which mark differentiating HSCs, showed a significant enrichment upon IR (**Figure S2C)**. Altogether, these data show that IR induces a loss of transcriptional signatures involved in HSC quiescence, long-term potency and self-renewal capacity, and a gain in gene signatures involved in HSC differentiation, thus recapitulating the HSC loss of self-renewal previously reported upon IR (de Laval et al., 2013).

### IR-induced downregulation of gene expression is associated with the presence of L1Md in their introns

In order to check if gene deregulation upon IR might be associated with H3K9me3 enrichment changes at gene promoters, we quantified H3K9me3 enrichment over promoter regions (−2kb; +1kb around TSS; U-MRA). We found 239 promoter sequences showing a significant (*p<0.05*) H3K9me3 differential enrichment upon IR, 211 and 28 showing decreased and increased H3K9me3, respectively **(Figure S2D)**. However, these variations are not correlated with gene upregulation, and only very poorly correlated with gene downregulation (**Figures 4A and S2E)**.

**Figure 4.**
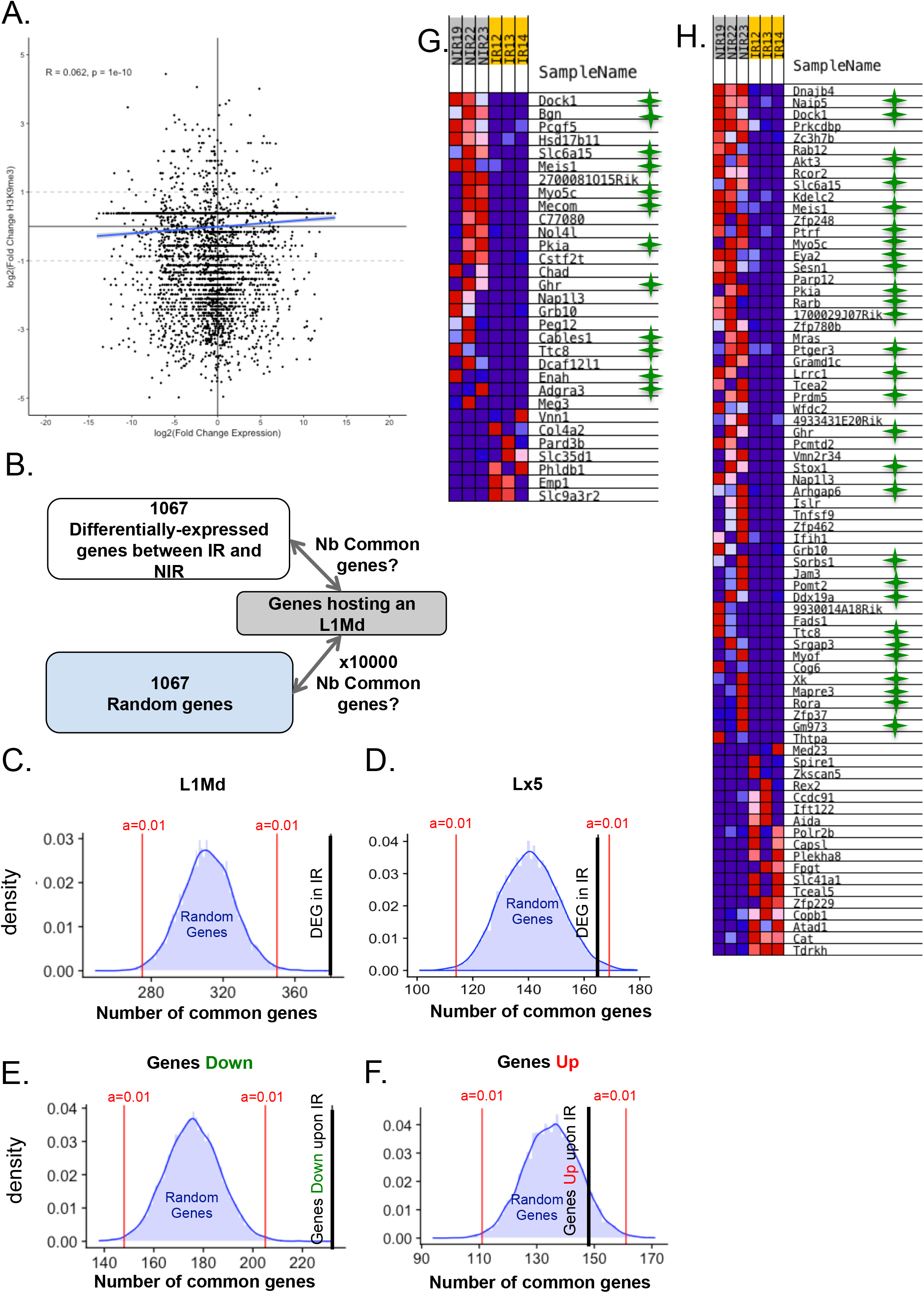
Gene repression upon IR is associated with intronic L1Md. **(A)** Correlation plot representing the log2(Fold change) in H3K9me3 concentration at gene promoters *vs* log2(Fold change) in gene expression. (**B-F)** Permutation test comparing of the number of genes found in common between the list of DEG or 10000 lists of random genes and the list of genes hosting one or several L1Md. Blue curve: Distribution of the number of genes found in common between random genes and genes hosting an L1. Black vertical line: number of genes found in common between DEG and genes hosting an L1Md (**C)** or an Lx5 (**D)**; or between genes downregulated (**E)** or upregulated (**F)** upon IR and hosting an L1Md. Significance bars (*p<0.01*) are shown in red. (**H-I)** Heatmaps of the expression of genes from two LT-HSC signatures that are significantly up- (red) or down- (blue) regulated in IR vs NIR. Green stars indicate the presence of an intronic L1Md in the downregulated gene.

Since the loss of H3K9me3 upon IR mainly occurs at L1Md, we sought to determine if L1Md derepression might be involved in the gene deregulation observed after IR. Most of the information concerning H3K9me3 enrichment at young RTE such as L1Md is obtained through M-MRA. However, reads from the M-MRA are arbitrary assigned and cannot be precisely localized. Thus it is impossible to determine if L1Md derepression is associated with gene deregulation by crossing our ChIP-seq and RNA-seq data. To overcome this issue, we crossed the list of the 1067 DEGs (*p<0 .05*) in IR *vs* NIR from our RNA-seq data with the list of genes hosting one or several L1Md (reconstructed repeatMasker database) (**Figure 4B; Table S3)**. The vast majority of these L1Md are located in introns (99%) **(Figure S2F)**. We found that 377 DEGs in IR host one or several L1Md, in majority located in introns. This is significantly (*p<0.0001*) more that one would expect by chance, as revealed by a permutation test using 10,000 lists of 1067 genes randomly extracted from the refseq database (**Figures 4B and 4C**). These results reveal a strong and significant association between gene deregulation upon IR and the presence of an intronic L1Md in these genes. This is specific for L1Md as no significant association was observed between DEGs and the presence of Lx5, an older LINE subfamily (**Figure 4D**). Surprisingly, this association is specific for genes that are downregulated upon IR (**Figure 4E**) and was not found for upregulated genes (**Figure 4F; Table S3)**.

Interestingly, 50% of the genes from LT-HSC signatures (Chambers et al., 2007; Pietras et al., 2015) whose expression is significantly decreased upon IR (**Figures 4G and 4H**) contain one or several L1Md in their introns. Similarly, we found that 31% and 41% of the genes from the low-output and MK-biased signatures (Rodriguez-Fraticelli et al., 2020) host one or several L1Md **(Figures S2G and S2H)**. Altogether, these data highlight a significant association between genes whose expression is repressed upon IR, notably those belonging to the HSC signature, and the presence of intronic L1Md.

### Gene repression upon IR is associated with loss of H3K9me3 at selected intronic L1Md loci harboring NF-κB binding sites

We next investigated if the loss of H3K9me3 induced by IR at intronic L1Md in a given gene could be associated with its decreased expression in *cis*. For this purpose we chose 3 HSC candidate genes whose expression is significantly reduced by IR in our RNA-seq data : *Mecom*, *Pkia* and *Ttc8* (**Table S2**), as well as 3 negative controls, also harboring at least one intronic L1Md but whose expression remains unchanged upon IR (0.7< FC <1.3, p>0.05): *Snx27*, *Mapre2*, and *Celf2*. One month after TBI, a significant downregulation of *Mecom*, *Pkia* and *Ttc8*, but not of the 3 controls was observed (**Figures 5A and 5B**). To assess H3K9me3 enrichment specifically at intronic L1Md of these genes, we performed H3K9me3 ChIP-qPCR experiments using primer pairs located both in the intron and in the 5’ end of the L1Md, thus allowing unique and specific amplicon production (**Figure 5C**). We chose in each case the longest L1Md (>5kb), displaying higher enrichment at the 5’ end (**Figure 2H),** to detect significant basal H3K9me3. We first confirmed that H3K9me3 is indeed present at all intronic L1Md, as compared to *Spi1* and 5S negative controls **(Figures S3A and S3B).** Of note, H3K9me3 levels at all these L1Md are similar to that found globally at L1Md_A promoters (**Figure S3B**). IR induced a significant decrease in H3K9me3 specifically at all the chosen canditate intronic L1Md of genes downregulated upon IR, but not at the control genes (**Figure 5D**). Of note, we randomly chose 3 negative candidates that fit our criteria, and for the 3 candidates selected, no loss of H3K9me3 was detected. This indicates that IR-induced H3K9me3 loss at intronic L1Md is not a general event but rather that it seems to occur only in specific genes whose expression is reduced upon IR. This suggests that the presence of the H3K9me3 mark at this location may play a role in the regulation of gene expression *in cis* upon IR.

**Figure 5.**
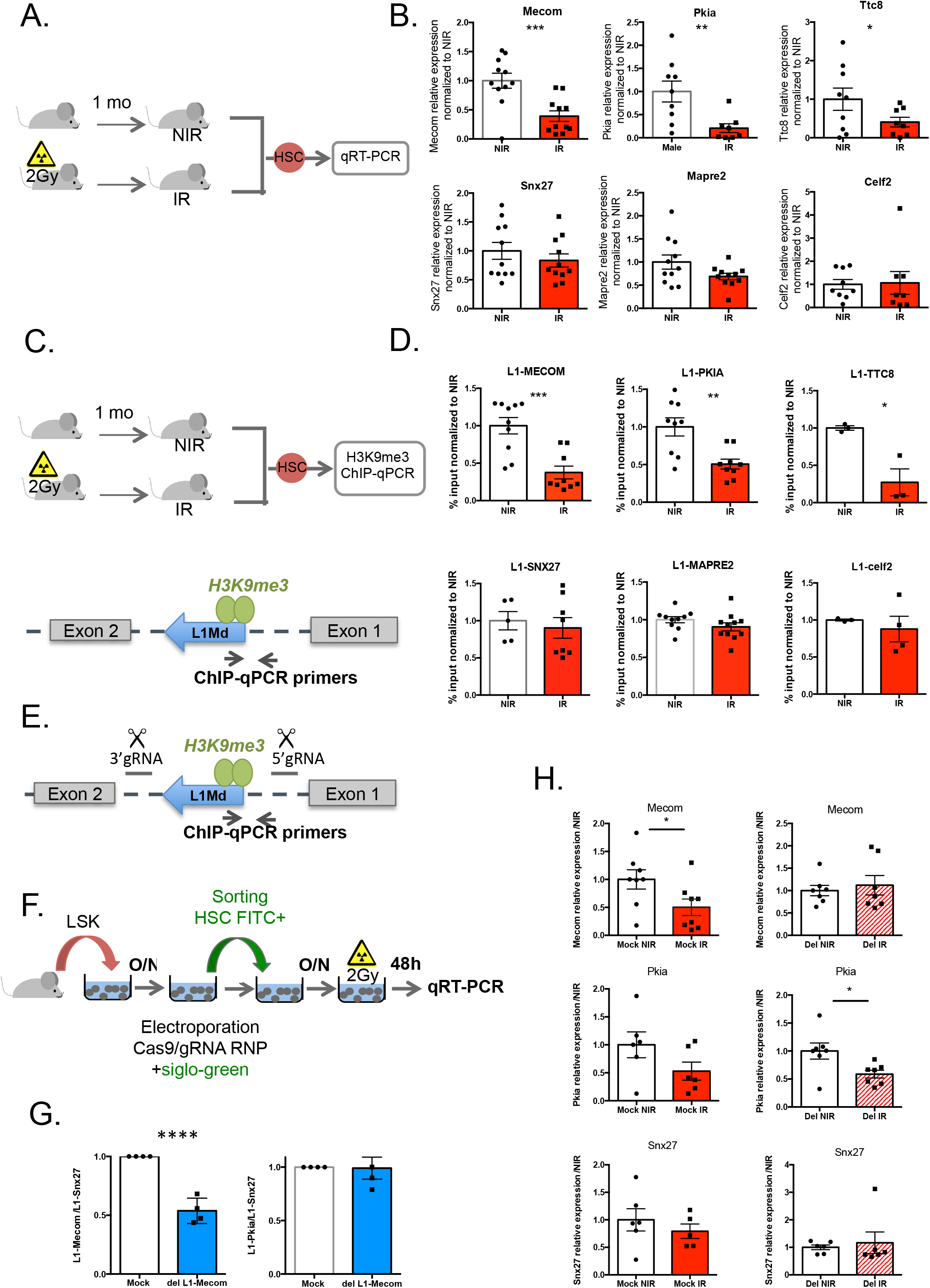
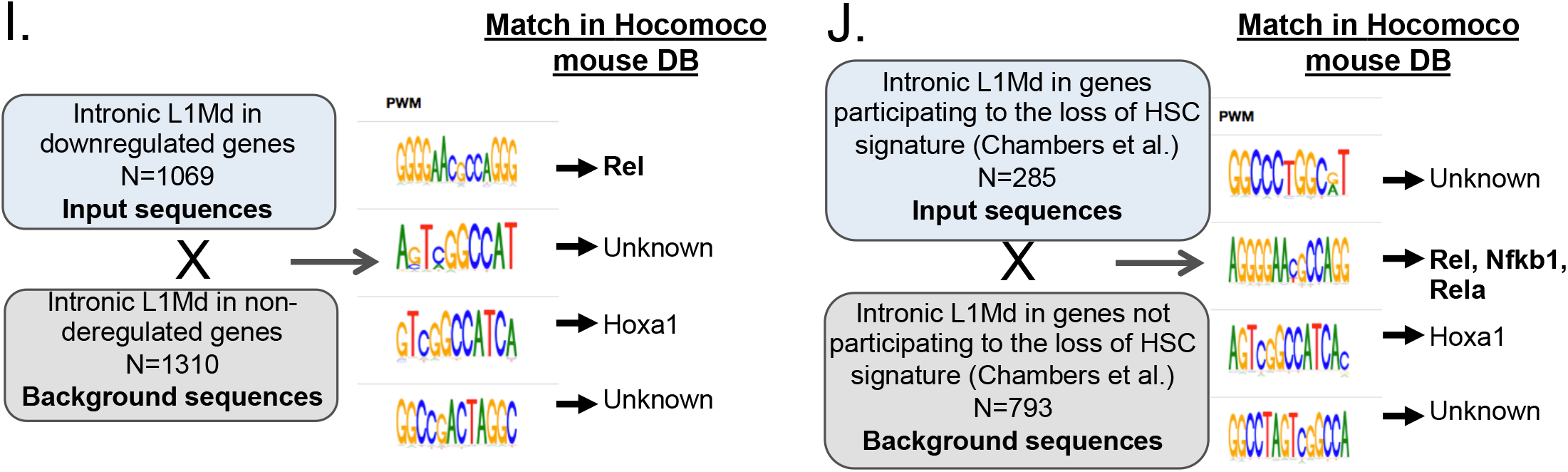
Gene repression upon IR is associated with loss of H3K9me3 at intronic L1Md loci harboring NF-κB binding sites. **(A, B)** Experimental design and mRNA expression assessed by RT-qPCR in HSCs one month post TBI. Ct values were normalized to RPL32 and HPRT. Results are expressed as fold change from the mean value of the NIR condition. Each dot represents a pool of 3 (NIR) or 4 (IR) mice. Means +/− SEM from 3 to 4 independent experiments. **(C, D)** Experimental design and H3K9me3 ChIP-qPCR enrichment 1 month post TBI. The primer positioning at the intronic L1Md allowing the amplification of a unique and specific product is shown (**C**). Each dot represents a pool of 3 (NIR) or 4 (IR) mice from 2-4 independent experiments. Results are means +/− SEM of the percentage of input normalized to the NIR condition. ***p<0.001; **p<0.01; *p<0.05 t-test. **(E-H)** CRISPR/Cas9-induced deletion intronic L1Md in *Mecom* gene. **(E)** gRNA positioning around L1Md sequence**, (F)** experimental design, **(G)** DNA amplification assessed by RT-qPCR and **(H)** mRNA expression assessed by RT-qPCR in Cas9-gRNA RNP electroporated HSCs 48h after irradiation *in vitro.* Ct values were normalized to L1-Snx27 for DNA amplification and to Rpl32 for mRNA quantification. mRNA expression was normalized to NIR values. Each dot represents a pool of electroporated HSCs from 3 independent experiments. *p<0.05 t-test. **(I-J)** *De novo* motif discovery analysis performed with the BaMMmotif tool on L1Md sequences located in introns of downregulated genes *vs* non-deregulated genes **(I)** or the genes participating *vs* not participating to the loss of the LT-HSC signature **(J)**. Enriched motifs were matched to known motifs using the Hocomoco mouse database.

In order to test this hypothesis, we chose to delete specifically the intronic L1Md of *Mecom* through CRISPR/Cas9 and guide RNAs targeting each side of the L1Md (5’gRNA and 3’gRNA) (**Figure 5E**) and tested *Mecom* expression upon IR. We electroporated LSK cells with the Cas9/gRNA RNP complex (Gundry et al., 2016) together with a siglo green electroporation indicator and we sorted electoprated (FITC+) HSCs for analysis (**Figures 5F and S3C**). We first tested different combinations of 5’ and 3’ gRNAs **(Table S4)**. Only one of these combinations was efficient in deleting the L1Md **(Figure S3D)**. Using this combination, we then validated that L1Md deletion occurred in *Mecom*, but not in *Pkia*, through qRT-PCR using the ChIP-qPCR primers (**Figure 5G**). We also validated that in mock electroporated cells IR could induce the specific downregulation of *Mecom* and *Pkia* but not of *Snx27* expression *in vitro*, as observed *in vivo* (**Figure 5H**). While *Pkia* expression remained significantly reduced upon IR after *Mecom* intronic L1Md deletion, *Mecom* expression was not affected anymore. This suggests that *Mecom* intronic L1Md acts in cis, and is responsible for the specific *gene* downregulation upon IR (**Figure 5H).**

To unravel what makes the specificity of H3K9me3 loss at L1Md upon IR, we interrogated the potential enrichment for binding motifs for transcription factors in the L1Md located in genes downregulated in IR (*p<0.05*, 1069 L1Md = input sequences) *vs* non-deregulated genes (*p>0.05*, 0.7<FC<1.3, 1310 L1Md = background sequences) through *de novo* motif search using BAMMmotif (https://bammmotif.soedinglab.org/) (Kiesel et al., 2018). Of the 4 motifs found significantly enriched in L1Md located in downregulated genes, 2 were not retrieved in the mouse Hocomoco motif database, one motif corresponds to the binding site of Hoxa1, and one to the binding site of Rel, a member of the NF-κB transcription factor family (**Figure 5I**). D*e novo* motif search in the L1Md located in downregulated genes *vs* upregulated genes also retrieved Rel motif (**Figure S3E**). Furthermore, *de novo* search for motifs specifically enriched in L1Md located in genes participating (284 L1Md) *vs* genes not participating (793 L1Md) to the loss of the HSC signature (**Figure 3H**) also revealed significant enrichment for NF-κB transcription factors Rel, RelA and Nfkb1 (**Figure 5J**). Similar results were found for the low-ouput / self-renewing LT-HSC signatures **(Figures S3F and S3G)**. Of note, this motif is present in the intronic L1Md of IR-regulated genes *Mecom*, *Pkia* and *Ttc8* which loose H3K9me3 upon IR (**Figure 5D, Table S3**), supporting the possibility that it may regulate the presence of H3K9me3 at these loci.

Performing *de novo* search on promoter sequences of genes downregulated by IR (p<0.05, 3893 promoter sequences) *vs* non-deregulated genes (0.7<FC<1.3, p>0.05, 14208 promoter sequences from which we randomly extracted 3893 sequences) did not show specific enrichment in motifs for NF-κB members. Instead, motifs for different transcription factors such as Egr, Znf, Foxj, Foxq were found **(Figure S3H)**.

Altogether, these data suggest that the NF-κB pathway may control HSC gene expression by regulating the presence of the H3K9me3 mark at L1Md located in introns, and not by affecting promoters.

### TNFα treatment prevents loss of H3H9me3 at intronic L1Md, HSC gene repression and HSC loss of function during IR stress

In mammals, the NF-κB family is composed of five members: RELA (p65), RELB, c-REL, and the precursor proteins NFKB1 (p105) and NFKB2 (p100), which are processed into their active forms, p50 and p52 respectively, and are active as homo or heterodimers (Cartwright et al., 2016). The canonical NF-κB pathway involves p50, p65 and c-Rel. P50 lacks transactivation domain. Thus, while p50/p65 or c-Rel heterodimers act as transcriptional activators, p50/p50 (NFKB1) homodimers are generally described as transcriptional repressors. NFKB1 notably represses the expression of proinflammatory genes through the recruitment of chromatin modifiers and H3K9 methylation (Ea et al., 2012; Elsharkawy et al., 2010). In addition, p50 has been shown to shuttle between the nucleus and cytoplasm and to bind to a large number of genes in unstimulated cells (Schreiber et al., 2006). This makes of this factor a good candidate to regulate H3K9me3 levels at L1Md presenting NF-κB sites and IR-induced HSC gene expression changes. Supporting this possibility, HSCs sorted from mice one month after TBI showed a significant decrease in both NFKB1 mRNA, as well as protein expression tested with two different antibodies (**Figures 6A-C and S4A**). Processing of p105 to p50 is regulated both independently of the NF-κB activation pathway and during activation of the canonical pathway induced by proinflammatory cytokines such as TNF-α (Cartwright et al., 2016). As shown above, IR induces a loss of the TNFA_signaling_Via_NFKB signature (**Figures 3D and 3E**). Thus, we asked whether TNF-α stimulation could prevent IR effects by rescuing the levels of p50 homodimers. NFKB1 protein expression decreased after 48h of culture of purified HSCs that have been irradiated *in vitro* (**Figures 6D**, **6E and S4B)**. Addition of TNF-α to the cell medium prior to IR prevented this effect. As after TBI, the loss of NFKB1 induced by IR *in vitro* was associated with a specific decrease of *Mecom*, *Pkia* and *Ttc8* but not of *Celf2*, *Snx27* and *Mapre2* mRNAs (**Figures 6F and S4C)**. Importantly, H3K9me3 ChIP-qPCR at intronic L1Md of the selected genes correlated with gene expression with a significant and specific reduction at *Mecom* and *Pkia* intronic L1Md but not at *Snx27* and *Celf2* (**Figure 6G**). This shows that IR-induced H3K9me3 loss at intronic L1Md and gene expression changes in HSCs are direct and short-term and that TNF-α stimulation can rescue both effects. Supporting a role of NFKB1 in these effects, *Nfkb1*^−/−^ HSCs showed reduced expression of *Mecom* and *Pkia* but not of *Snx27* mRNA when compared to wild-type (WT) HSCs (**Figure 6H**). In addition, TNF-α was unable to rescue HSC gene expression or H3K9me3 levels at their intronic L1Md in *Nfkb1*^−/−^ HSCs (**Figure 6I and J**).

**Figure 6.**
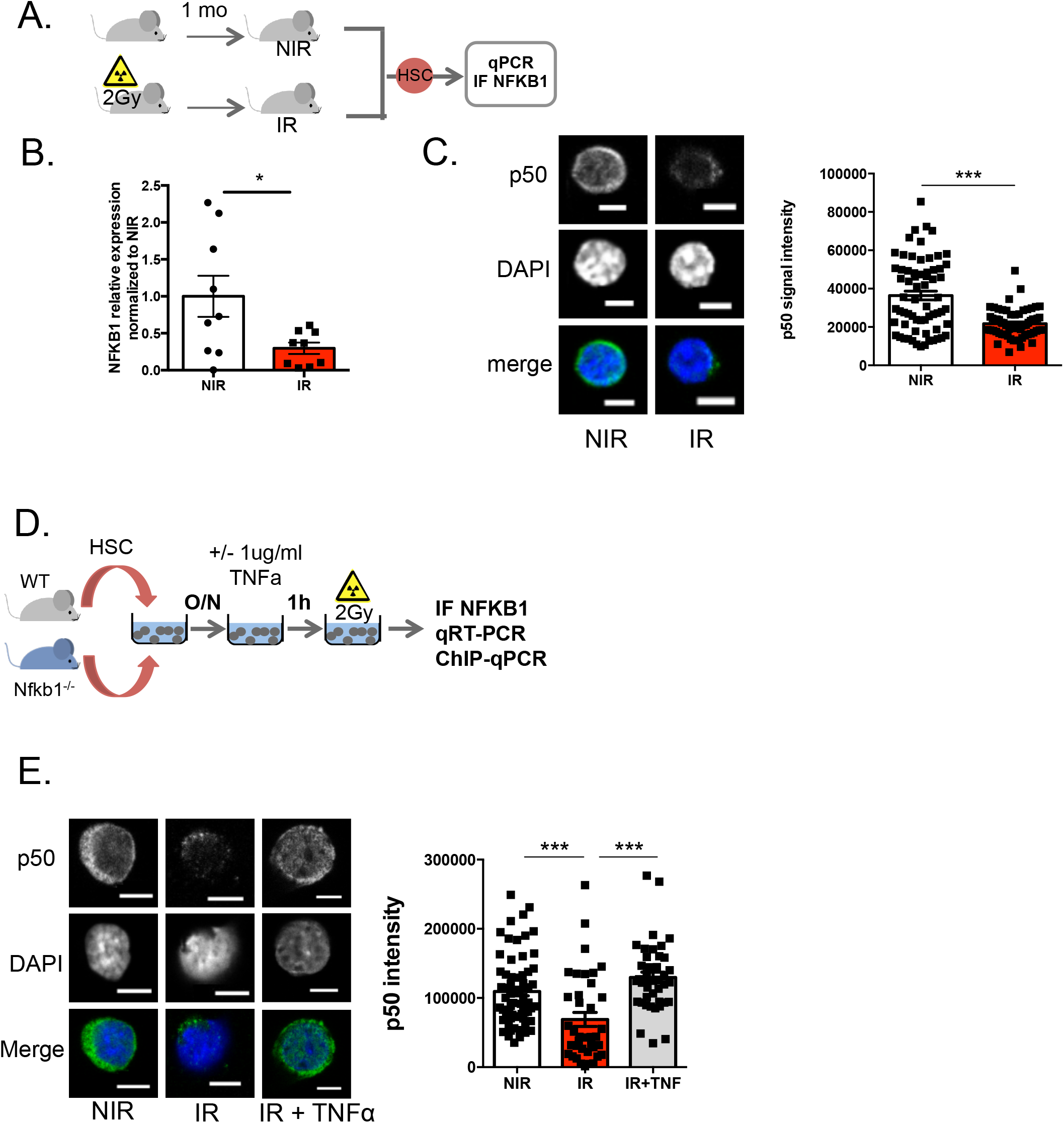
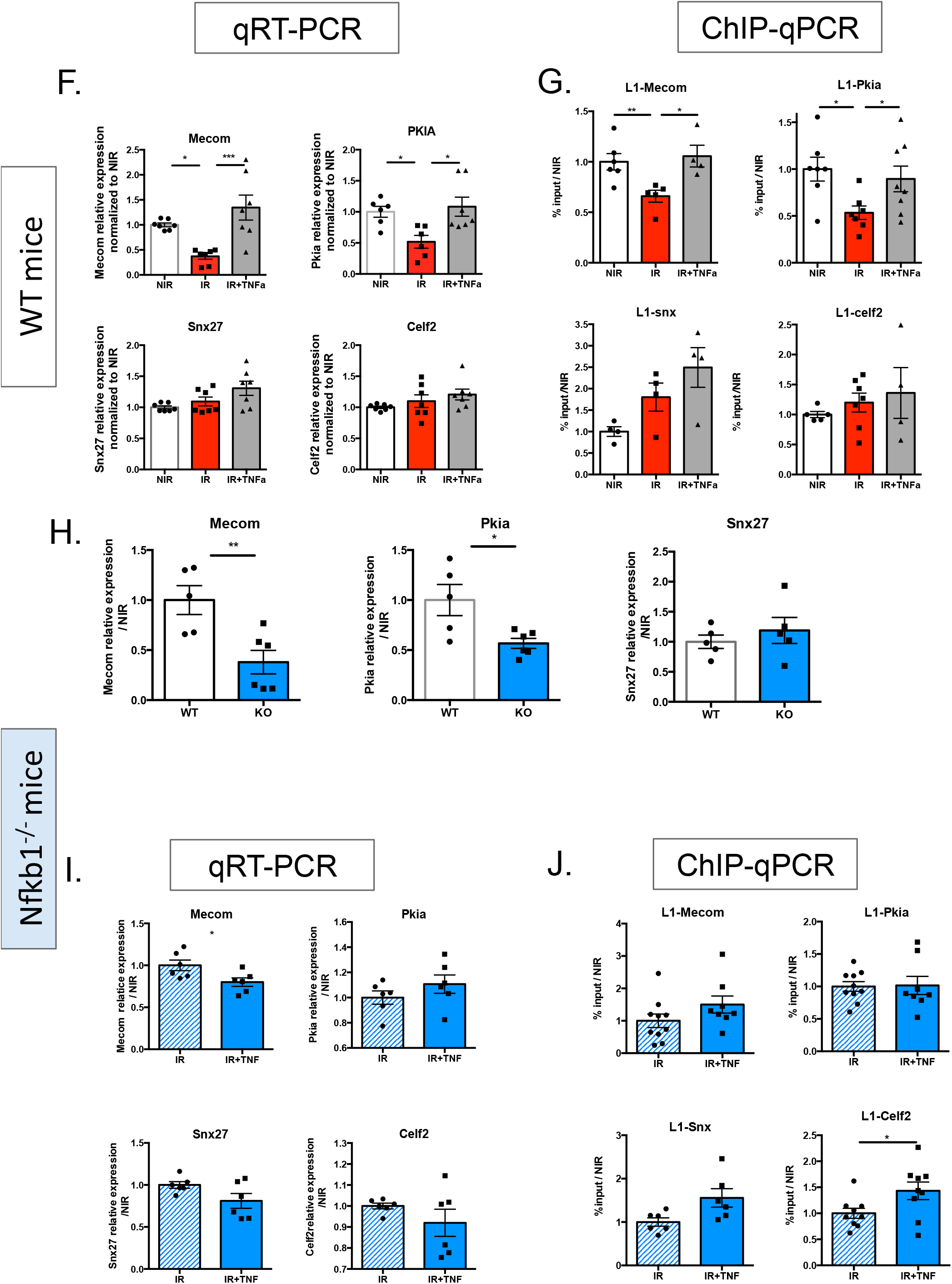
TNF-α treatment prevents loss of H3H9me3 at intronic L1Md and HSC gene repression *in vitro*. **(A-C)** NFKB1 expression in HSCs one month after TBI. (**B**) mRNA expression measured by RT-qPCR. Ct values were normalized to mean of RPL32 and HPRT. Results are expressed as fold change from the mean value of the NIR condition. Means +/− SEM. Each dot represents a pool of 3 (NIR) or 4 (IR) mice from 3 independent experiments. t-test**. (C)** Representative images and quantification of Nfkb1 protein mean IF intensity. Bars, 5µM. Each dot represents a cell. Results are expressed as fold change from the mean value of the NIR condition from 2 independent experiments and represented as means +/− SEM. t-test. **(D)** Experimental design analyzing the effects of IR and TNF-α *in vitro* in WT and *Nfkb1*^−/−^ mice. **(E)** Representative images and quantification of Nfkb1 staining. Bars, 5µM. Each dot represents a cell. Results are represented as mean +/− SEM of Nfkb1 IF intensity. One-way ANOVA with Tukey’s multiple comparison test. **(F)** Gene expression evaluated by RT-qPCR in WT mice. Means +/− SEM from 2 independent experiments. One-way ANOVA with Tukey’s multiple comparison test. **(G)** H3K9me3 enrichment at intronic L1Md evaluated by ChIP-qPCR. Results are expressed as in legend to Figure 5B. Means +/− SEM from 2-3 independent experiments. One-way ANOVA with Bonferroni’s multiple comparison test. **(H)** Experimental design for reconstitution experiments using HSC sorted from mice one month after TBI and previously treated with TNF-α (IR+TNF) or not (IR), or left untreated (NIR); BMMC, bone marrow mononuclear cells. **(I-J)** percentage of GFP-negative donor contribution in blood in mice transplanted with NIR, IR or IR+TNF cells at 7 **(I)** and 14 weeks **(J)** after reconstitution. **(K, L)** LSK CD34^−^Flk2^−^GFP^−^ **(K)** or LSK CD34^−^Flk2^−^CD48^−^ **(L)** GFP-negative donor HSC contribution in the BM 14 weeks after reconstitution. One-way ANOVA Tukey’s multiple comparison test. *p<0.05; **p<0.01; *** p<0.001; ****p<0.0001. **(F)** Model. At basal, the NFKB pathway, possibly through its repressor NFKB1 (p50/p50 homodimers), is involved in the recruitment of H3K9 methylases (HMT) at intronic L1Md enriched in NFKB binding sites motifs, and apposition of the repressive histone mark H3K9me3. H3K9me3 “islands” into the body of transcribed genes may help the processing of RNAPolymerase II (RNAPolII) and transcript stability. Upon Irradiation, loss of the TNF-α-NFKB pathway leads to a loss of H3K9me3 at the intronic L1Md, gene repression and transcript stability. Alternatively, loss of H3K9me3 at intronic L1Md may lead to their transcription and gene repression through transcriptional interference.

We then investigated if TNF-α stimulation could prevent IR effects *in vivo.* WT mice received 2 injections with 2ug TNF-α at 12h interval and were irradiated one hour after the last injection. We then performed H3K9me3 Cut&Tag and RNA-seq in HSCs sorted one month after IR (**Figure 7A).** Cut&Tag gave an efficient profiling of H3K9me3 with exceptionally low background as observed at the SUV39h1 and 2-dependent H3K9me3 peaks (Bulut-Karslioglu et al., 2014) (**Figure S5A**) and as previously described (Kaya-Okur et al., 2019). Plot profile of H3K9me3 enrichment along L1Md sequences confirmed the loss of H3K9me3 at L1Md upon IR and showed it could be prevented by TNF-α (**Figure 7B**). Notably, TNF-α also inhibited the specific loss of H3K9me3 enrichment at the intronic L1Md of *Mecom* and *Pkia* but not *Mapre2* (**Figure 7C**) and restores the corresponding gene expression (**Figure 7D**). RNA-seq analysis further showed that TNF-α injection *in vivo* prevented IR-induced loss of both TNF-α *via* NF-κB (**Figure 7E**) and long-term HSC signatures (**Figure 7F and S5B.)**.

**Figure 7.**
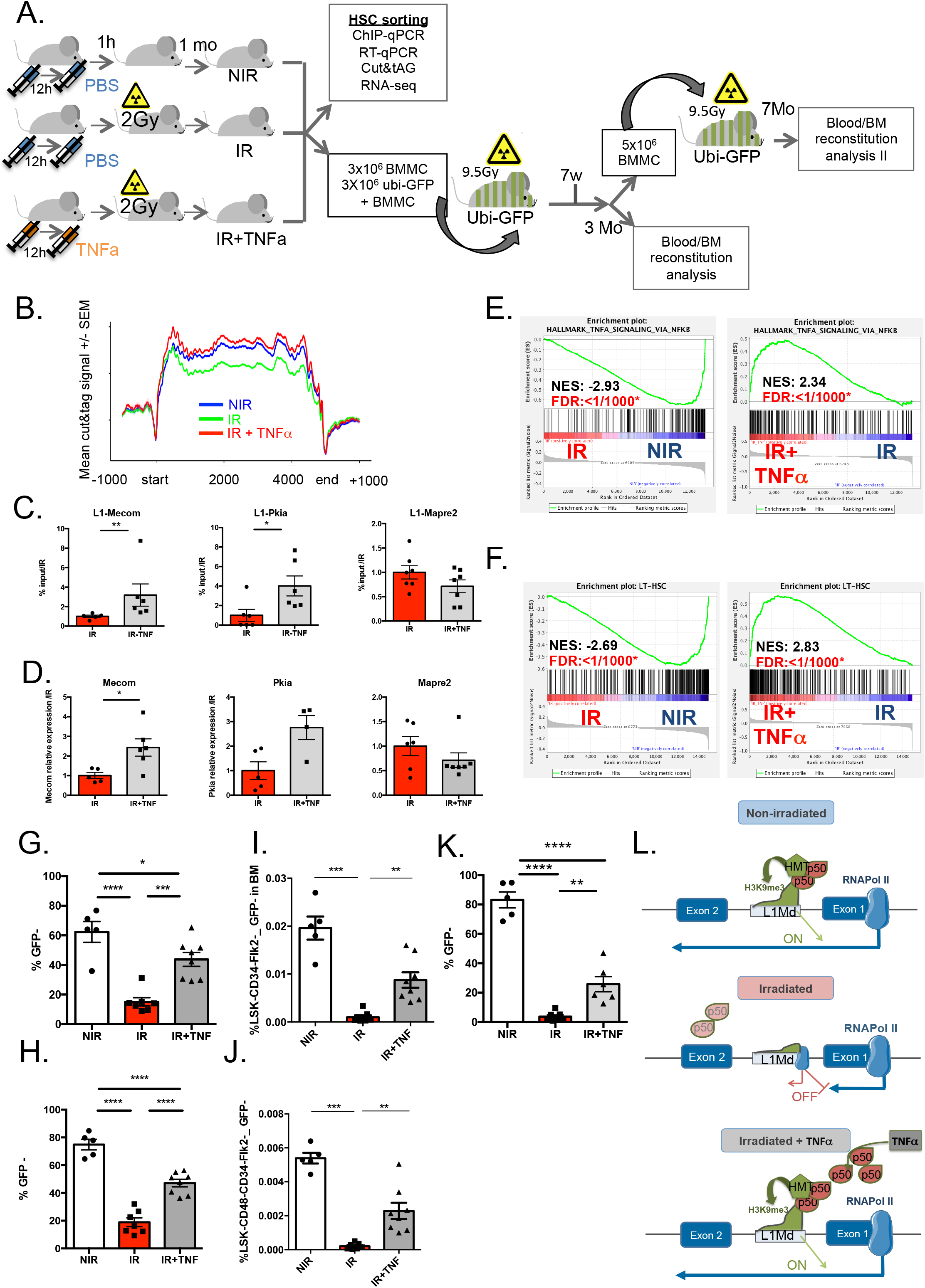
TNF-α treatment prevents loss of H3H9me3 at intronic L1Md, HSC gene repression and HSC function *in vivo*. **(A)** Experimental design for TNF-α treatment *in vivo,* molecular analysis and reconstitution experiments using HSCs sorted from mice one month after TBI and previously treated with TNF-α (IR+TNF) or not (IR), or left untreated (NIR); BMMC, bone marrow mononuclear cells. **(B)** Plot profile of H3K9me3 enrichment along the L1Md sequences +/− 1kb flanking regions in NIR (blue), IR (green) and IR+ TNF-α (red) conditions. Each line represent the merged Cut&Tag signal from 2 (NIR) to 3 (IR and IR+ TNF-α) mice +/− SEM. **(C)** H3K9me3 enrichment at intronic L1Md evaluated by ChIP-qPCR. **(D.)** mRNA expression measured by RT-qPCR. Ct values were normalized to RPL32 and HPRT. Results are expressed as fold change from the mean value of the IR condition. Each dot represents a pool of 4 mice. Means +/− SEM from 2 independent experiments, t-test. **(E-F)** GSEA analysis using Gene Sets. Enrichment plots for TNF-α_signaling_via_NFKB hallmark (**E),** and LT-HSC signature (**F). (G-H)** percentage of GFP-negative donor contribution in blood in mice transplanted with NIR, IR or IR+TNF cells at 7 **(G)** and 14 weeks **(H)** after reconstitution. **(I,J)** LSK CD34^−^Flk2^−^GFP^−^ **(I)** or LSK CD34^−^Flk2^−^CD48^−^ **(J)** GFP-negative donor HSC contribution in the BM 14 weeks after reconstitution. One-way ANOVA Tukey’s multiple comparison test **(K)** percentage of GFP-negative donor contribution in blood in mice secondary transplanted with pool of NIR, IR or IR+TNF mice from the primary reconstitution. One-way ANOVA Tukey’s multiple comparison test. (**L**) Model. At basal, the NF-κB pathway, possibly through its repressor NFKB1 (p50/p50 homodimers), is involved in the recruitment of H3K9 methylases (HMT) at intronic L1Md enriched in NFKB binding sites motifs, and apposition of the repressive histone mark H3K9me3. H3K9me3 “islands” into the body of transcribed genes may help the processing of RNAPolymerase II (RNAPolII) and transcript stability. Upon Irradiation, loss of the TNF-α-NF-κB pathway leads to a loss of H3K9me3 at the intronic L1Md, gene repression and transcript stability. This is prevented by prior TNF-α treatment. *p<0.05; **p<0.01; *** p<0.001; ****p<0.0001.

Finally, to confirm the importance of this pathway in HSC maintenance, we analyzed the effect of TNF-α treatment on HSC function upon IR. Total BM cells isolated from mice treated with TNF-α before TBI or not were transplanted in competition with total BM cells from mice ubiquitously expressing GFP (ubi-GFP mice) into lethally irradiated ubi-GFP mice (Schaefer et al., 2001) (**Figure 7A**). Three months after reconstitution, as expected, the percentage of GFP negative IR donnor cells in the blood was greatly decreased. TNF-α treatment before TBI significantly prevented this effect (**Figures 7G, H and S5C**). It also significantly prevented IR-induced LT-HSC loss (**Figures 7I, J and S5D).** Secondary transplants showed that the self-renewal function of HSCs after IR could be restored by TNF-α treatment (**Figure 7K).**

Altogether, these data suggest that TNF-α treatment rescues HSC reconstitution ability upon IR by preventing IR-induced decrease in NFKB1 repressor expression, specific derepression of L1Md harboring NF-κB binding sites in the introns of HSC genes, and thereby their repression (**Figure 7L**).

## DISCUSSION

Epigenetic factors controlling DNA and histone methylation are key regulators of HSC function and are often mutated in leukemia. These mutations also increase with age in otherwise healthy individuals where they confer clonal hematopoiesis. However, epigenetic alterations have been observed in aged HSCs in the absence of mutations in epigenetic factors (Djeghloul et al., 2016; Sun et al., 2014). HSC aging phenotype is reversible, suggesting that it results from an altered epigenome (Wahlestedt et al., 2017).

H3K9me3 alterations is a hallmark of aging and cellular senescence in model organisms (Criscione et al., 2016; Ocampo et al., 2016). Although H3K9me3 has been shown to be crucial for HSC identity (Koide et al., 2016), H3K9me3 changes in HSCs have, to our knowledge, never been studied in the context of stress such as IR. We show here that IR stress profoundly affect HSC heterochromatin by significantly reducing H3K9me3 enrichment, without affecting H3K9 methyltransferase (SETDB1, KAP1, or MPP8 from the HUSH complex,) and that this loss mainly occurs at L1Md elements. This effect was observed both after short time *in vitro* and long time after TBI *in vivo*, suggesting that heterochromatin alterations may explain the long-term effect of IR on HSC function.

Heterochromatin alterations are associated with a strong deregulation of the HSC transcriptome. H3K9me3 enrichment at promoters has recently emerged as a key player in the repression of lineage-inappropriate genes (Koide et al., 2016). Surprisingly, we found here that gene deregulation is not associated with H3K9me3 changes at gene promoters, but is rather associated with the loss of H3K9me3 at intronic L1Md. H3K9me3 enriched at intronic evolutionary recent L1Md was previously shown to be involved in the tight regulation of gene transcription in ESCs (Liu et al., 2018). Some ERVs also play the role of AML enhancers with a driving role in leukemia cell phenotype (Deniz et al., 2020). However, our study is the first showing the involvement of L1Md on the regulation of HSC gene expression.

With CRISPR/Cas9 targeted deletion, we show that the presence of intronic L1Md in *Mecom* is required for *Mecom* gene specific downregulation upon IR and suggests that intronic L1Md can act in cis to regulate gene expression. Although surprising at first glance, repression of genes following derepression of intragenic L1 was previously reported in cancers (Aporntewan et al., 2011). This may be due to transcriptional interference (Han et al., 2004; Kaer et al., 2011; Ninova et al., 2020), mostly through intron retention, exonization and cryptic polyAdenylation (Kaer et al., 2011). H3K9me3 is mostly enriched in the 5’UTR part of the L1Md. Loss of this repressive mark at the L1Md promoter regions may induce its transcription and gene repression through transcriptional interference. We cannot test directly this hypothesis due to the repetitive nature of these sequences. However, gene repression is associated with the presence of L1Md in their introns whatever their size, suggesting that gene repression may not be exclusively due to intronic L1Md transcription.

Instead of blocking transcription, DNA methylation or H3K9me3 within gene bodies is a feature of transcribed gene (Jones, 2012; Ninova et al., 2020; Vakoc et al., 2005). H3K9me3 loss in gene bodies was previously shown to be associated with gene repression (Ninova et al., 2020). The presence of H3K9me3 islands in the body of the genes has been proposed to slow-down RNA Polymerase II (RNAP II) elongation rate (Saint-André et al., 2011; Vakoc et al., 2005). This was shown to help the recognition of true *vs* cryptic RNA processing sites, controlling alternative splicing (de la Mata et al., 2003)(De La Mata et al., 2003)(De La Mata et al., 2003)(De La Mata et al., 2003), polyadenylation and finally transcript stability. This is particularly relevant in the case of genes bearing long introns. Interestingly, the genes downregulated upon IR in HSCs are significantly longer than by chance (data not shown). Slowing down RNAP II elongation rate in these long intron genes might also prevent R-loop formation and genomic instability (Aguilera and Gaillard, 2014). Interestingly, evolutionary young L1 located in long introns recruit RNA-binding proteins to prevent improper splicing or polyAdenylation (Attig et al., 2018). Thus, even in the absence of intronic L1Md transcription, loss of H3K9me3 islands enriched at intronic L1Md in the body of the genes might be involved in splicing defects and/or premature polyadenylation. Whether these mechanisms are involved in HSC gene regulation will require further investigations.

Our RNA-seq analysis indicates a strong downregulation of the TNF-α-NF-κB gene signature one month after TBI. IR specifically reduced the recently described HSC-prosurvival TNFα- NF-κB signature required to maintain HSCs during inflammation or cytotoxic BM ablation (Yamashita and Passegué, 2019). This suggests that loss of long-term regenerating HSCs and TNF-α-NF-κB gene expression upon IR may be linked. Supporting this possibility, treatment of HSCs with TNF-α before IR *in vitro*, or its injection to mice before TBI, restored HSC gene expression and their reconstitution ability. These results strongly support previous data showing that TNF-α-NF-κB signaling is required to regulate HSC function under stress (Hu et al., 2021; de Laval et al., 2014; Yamashita and Passegué, 2019). TNF-α promotes HSC survival through p65/RelA NF-κB subunit (Yamashita and Passegué, 2019). This factor has also been found to control the expression of genes involved in HSC maintenance (Stein and Baldwin, 2013). However, the promoters of HSC genes downregulated upon IR are not enriched in NF-κB binding sites. We show that L1Md associated with gene repressed upon IR are specifically and significantly enriched in NF-κB binding sites, and that this pathway regulates gene expression by controlling the level of H3K9me3 at these sequences. Interestingly, H3K9me3 enriched genomic regions specific to human ESCs as compared to more differentiated cells are enriched in NF-κB binding sites, suggesting their importance in establishing and maintaining the pluripotent state (Whitaker et al., 2015).

Whereas most of the NF-κB members can form active transcription factors, NFKB1 p50 subunit lacks transactivation domain and p50:p50 homodimers have been shown to act as stimulus-specific repressors notably during the resolution phase of inflammation, by recruiting H3K9 methyltransferases and HDAC at both NF-κB and type-I interferon (IFN) response genes (Cartwright et al., 2018; Ea et al., 2012; Elsharkawy et al., 2010). p50:p65 heterodimers are the most abundant form of NF-κB generated upon inflammatory stimuli. By contrast, p50 homodimers predominate in unstimulated cells where they can be prebound to the chromatin (Cartwright et al., 2016; Schreiber et al., 2006) suggesting that this factor may also play a role under non-inflammatory conditions. Indeed, we found that NFKB1 is present in both the cytoplasm and the nucleus of resting HSCs. Deletion of *Nfkb1,* or downregulation of p50/NFKB1 gene and protein as induced by IR, correlates with decrease of both gene expression and H3K9me3 levels at their intronic L1Md harboring an NF-κB binding sites. Conversely, increasing p50 production upon TNF-α stimulation rescued H3K9me3 levels at the specific intronic L1Md and gene expression in WT but not in *Nfkb1^−/−^* HSCs. This strongly supports the possibility that p50:p50 homodimers are the active repressor promoting the enrichment of H3K9me3 at L1Md located in HSC genes and the *cis-*regulation the host gene.

A growing body of evidence indicates that TEs have been coopted for transcriptional regulation in different cell and tissue types (Chuong et al., 2017). TEs are reservoirs of functional transcription factor binding sites. Since these sequence are widespread in the genome, they are largely contributing to the innovation of regulatory networks in a tissue-specific fashion (Chuong et al., 2017; Hermant and Torres-Padilla, 2021; Sundaram and Wang, 2018). Although LTR dominate this relationship, search for binding motifs in young L1 in human and mouse has revealed the presence of various TF motifs, including CTCF, YY1 and MYC (Sun et al., 2018; Sundaram and Wang, 2018). Our results show that NF-κB motifs are specifically enriched in most of the intronic L1Md sequences of genes downregulated during IR stress, and involve as much as 96% of these genes (**Table S3**). The presence of NF-κB binding sites in TEs is reminiscent of a study reporting that, in the human genome, 11% of NF-κB-binding sites reside in specific Alu SINEs, and that the vast majority of sites bound by NF-κB do not correlate with changes in gene expression (Antonaki et al., 2011). Although it is not known how many of these NF-κB motifs present in intronic L1Md have a functional role, the ability of TNF-α to restore both H3K9me3 levels at the L1Md and the expression specifically of genes including NF-κB motif-enriched intronic L1Mds, strongly suggests that at least some of these NF-κB-TE associations can influence gene expression.

TEs have rewired the antiviral gene regulatory network and they have been shown to play a key role in the regulatory evolution of immune response. Strong but opposing forces are driving the coevolution of TEs and antiviral defence (Chuong et al., 2016; Moelling and Broecker, 2019). Many IFN / NF-κB-target genes are viral restriction factors and contribute to the immune control of both endogenous (i.e. TEs) and exogenous genomic parasites (Gazquez-Gutierrez et al., 2021; Schneider et al., 2014). We and others have previously shown that IFN-I signaling controls young L1Md expression and L1 retrotransposition in HSCs and various tissues (Barbieri et al., 2018; Goodier et al., 2015; Yu et al., 2015). However, through the formation of dsRNA or cytoplasmic cDNA resembling viral nucleic acids, TEs are sensed by the cells as invading viruses and promote the activation of IRF3 and NF-κB transcription factors and the major antiviral immune pathways (Gazquez-Gutierrez et al., 2021; Volkman and Stetson, 2014). Notably, TE-derived dsRNAs have been shown to provide the inflammatory signal necessary for HSC generation during embryonic development (Lefkopoulos et al., 2020). Intringinly, beside HSC maintenance genes, many genes involved in IFN and NF-κB immune response pathway are found among genes downregulated in IR presenting an intronic L1Md with a NF-κB binding site (**Table S3)**. These include IFN and NF-κB target genes known to control L1 retrotransposition and / or levels such as *EiF2ak2* (Interferon-Inducible protein kinase R) and *Oas1g* RNase L whose activity are triggered by virus- or TE-derived dsRNAs ; *Jak2*, *Tyk2*, *Tnfrsf9* and *Birc2* involved in IFN and TNF responses, respectively, as well as T cell suppressing activity genes, *CD274* (PD-L1) and *CD86*. This further reinforces the causal relationships between TEs and immune genes and their coevolution. Interestingly, a higher TE occurrence has been found in immune gene-associated genomic regions and young TEs are specifically enriched in blood cells, as compared to other tissues (Trizzino et al., 2018; Ye et al., 2020). Enhancers unique for immune tissues are more prone to TE cooption, as compared to enhancers specific to other tissue types and the majority of them is conserved in HSC accessible chromatin regions. Using BAMMmotif for *de novo* motif search, we have found that the NF-κB motif is specifically enriched in L1Md that are present in genes of the HSC signature, and in the myeloid-leucocyte-mediated-immunity signature (GO:0002444) as compared to genes enriched in pancreas, testis, kidney, liver, placenta, salivary gland (Su et al., 2002) **(Figure S3I)**, or in genes from the immune system process (GO :0002376) as compared to genes from the reproductive process (GO :0022414) (**Figure S3J**). This suggests that NF-κB binding sites in L1Md might have been actively selected in introns of key HSC genes because of the immune-linked maintenance. This regulation might be important to expand the NF-κB and TNF-α activity by engaging more genes, including HSC maintenance genes into the NF-κB regulatory networks. Such activity could be important to maintain of HSCs while allowing expression of immune gene during TNF-α-NF-κB-induced myeloid regeneration or damage-induced bone marrow ablation, and further highlight the complex role of inflammation-induced pathways in HSCs. TNF-α levels are increased in patients with hematopoietic malignancies and the HSC-specific TNF-α signature is upregulated in MDS/AML malignant HSCs (Yamashita and Passegué, 2019). Exploring the mechanisms controlling TE expression and how inflammatory signals and aging impact them in normal and malignant HSC could lead to the identification of new selective dependencies of AML and new treatment strategies.

## METHODS

### Mice strains and treatments

Wild type (WT) C57BL/6J mice (6-8 week-old) were from the Envigo Laboratories. *Nfkb1*^−/−^ mice were from The Jackson Laboratory (B6.Cg-Nfkb1^tm1Bal^/J; Stock No:006097). All the mice were housed in a specific pathogen-free environment. All procedures were reviewed and approved by the Animal Care Committee (CE #2019_078_23286). Mice were injected retro-orbitally with 2µg TNF-α (Biolegend-Ozyme) before sublethal TBI (2 Gy) (RX irradiator X-RAD 320).

### Cell harvest and culture

Bone marrow was harvested from femur, tibia and hip bones in mice. Total bone marrow was depleted of differentiated hematopoietic cells (lineage-positive cells) using Mouse Hematopoietic Progenitor (Stem) Cell Enrichment Set (BD). Magnetically sorted Lineage-negative (lin^−^) cells were kept overnight at 4°C in IMDM medium supplemented with 10% FBS (HyClone) and 1% penicillin-streptomycin (Thermofisher). Staining was performed for 20min at room temperature (RT) using CD3ε (Lin) – APC clone 145-2C11 (553066, BD), TER-119 (Lin) – APC clone Ter-119 (557909, BD), CD45R/B220 (Lin) – APC clone RA3-6B2 (553092, BD), Ly6G-6C (Lin)-APC clone RB6-8C5 (553129, BD), Ly-6A/E (Sca-1) - PeCy7 clone d7 (558162, BD), CD117 (c-Kit) – PE or PerCP-Cy5.5, clone 2B8 (553355 or 560557 respectively, BD), CD34– FITC or AF700 clone RAM34 (560238 or 560518, BD), CD135 (Flk2) – BV421 or PE clone A2F10.1 (562898 or 553842 respectively, BD). HSCs (Lin^−^Sca^+^c-Kit^+^CD34^low^Flk2^−^) were sorted using ARIA3, ARIA Fusion or Influx cell sorters (BD Franklin Lakes, NJ, USA) and collected in Stem Span (StemCell).

When the cells were irradiated *in vitro*, HSCs were cultured in medium containing Flt3-Ligand, IL-3, IL-6, SCF, as described (de Laval et al., 2013) in the presence or absence of TNF-α. TNF-α was added to the medium at 1µg/ml 1 hour before IR.

### CRISPR-Cas9 deletion

Guide RNAs (gRNAs) were designed to generate specific deletion of the intronic L1Md of Mecom using CRISPOR (http://crispor.tefor.net/) (**Table S4**). 1ug total gRNAs (0.5ug 5’- gRNA+ 0.5ug 3’-gRNA) (Dharmacon) were incubated with 1ug Cas9 (CAS12205**-**Dharmacon) during 15min at RT and the Cas9-gRNA RNP was then co-electroporated with an equimolar siglo-green transfection indicator (D-001630-01-05) in 100 000 LSK after an O/N culture in medium containing Flt3-Ligand, IL-3, IL-6, SCF, as described (de Laval et al., 2013), and using a Neon transfection system (ThermoFisher Scientific) with the optimized electroporation condition 1700V, 20ms, 1pulse as previously described (Gundry et al., 2016). Just after electroporation, FITC+ HSC were sorted and collected in Stem Span (StemCell) containing Flt3-Ligand, IL-3, IL-6, SCF and left O/N in culture before irradiation. Cells were finally collected 48h after irradiation for further experiments.

### DNA extraction and genomic deletion verification

DNA from electroporated HSC was extracted using the tissue XS kit (Macherey-Nagel) according to manufacturer’s instructions, and the specific deletion of Mecom intronic L1Md was checked through qRT-PCR using the ChIP-qPCR primers (**Table S4**).

### Quantitative RT-PCR

HSCs were lysed in Tri-Reagent (Zymo Research) and stored at −80°C until used. Total RNA was extracted using the Direct-Zol RNA microprep kit (Zymo research) and reverse-transcribed with EZ Dnase VILO (Invitrogen). Real-time PCR was performed using the SYBR pPCR premix Ex Taq (Takara) or LUNA Universal qPCR Master Mix (NEB) on a 7500 real-time PCR machine (Applied Biosystems). Samples were tested for qPCR before reverse transcription to rule out detection of contaminating DNA. qPCR primers used were designed in different exons so as to minimize possible gDNA amplification. All data were normalized to the mean expression of RPL32, and HPRT. Primer sequences are shown in **Table S4**.

When necessary, 1.25µl of cDNA was preamplified for 14 PCR cycles in a multiplex reaction using Preamp Master-Mix (100-5580 - Fluidigm) and primer mix (200µM of each primer). To rule out primer dimerization or hairpin formation in the preamplification mix, primer sequences were previously analyzed using MFE3.0 PCR Primer Quality Control Software (Wang et al., 2019).

### ChIP-qPCR

10,000 HSCs were harvested in 1ml IMDM medium supplemented with 10% FBS and cross-linked using 1% formaldehyde (Invitrogen) for 10 min at RT. ChIP-qPCR experiments were performed using the True Micro-ChIP Kit (Diagenode) according to manufacturer’s instructions. Cells were sonicated using the Bioruptor Pico (Diagenode) sonication device for 10 cycles (20s ON/40s OFF). Chromatin was incubated overnight at 4°C using 0.25µg of H3K9me3 (C15410193-Diagenode) per IP. ChIP DNA was eluted and purified using the MicroChIP Diapure Columns (Diagenode). Subsequent qPCR Real-time PCR was performed as above ChIP-qPCR primers for intronic L1Md were designed such that one primer is located in the 5’ region of the L1Md, and the other primer is located in the intron of the host gene to allow the amplification of unique and specific product **(Table S4).**

### Immunofluorescence

3000-5000 HSCs were cytospun on glass slides and immunofluorescence was performed as previously described (de Laval et al., 2013). Two different Monoclonal anti-NFKB1 (p50) antibodies were used at 1/200: clone E10 was purchased from Santa Cruz Biotechnology, and clone D4P4D from Cell signalling. Detection was performed using Alexa Fluor 488-coupled anti-mouse secondary antibody (1/600). All slides were visualized using SPE confocal microscope (Leica). Pictures were analyzed using CellProfiler.

### Statistical analysis

Results were statistically evaluated using either the one-way ANOVA or unpaired t-test using GraphPad PrismTM version 6.0 software (GraphPad Software Inc., San Diego, CA, USA). The results are displayed as the means and SEM. The value of *p<0.05 was considered as significant, and **p<0.01 or ***p<0.001 as highly significant.

### RNA-seq and ChIP-seq reads quality

Quality of RNA-seq reads was assessed with Fastqc v0.11.8, Fastq-screen (Wingett and Andrews, 2018) v0.13.0 and MultiQC (Ewels et al., 2016) v1.7.

### RNA-seq

#### RNA quantification

Salmon (Patro et al., 2017) tool v0.14.1 was used to quantify mm10 NCBI RNA reference sequences (O’Leary et al., 2016) (RefSeq Curated, last updated 2017-11-16) downloaded from UCSC Table Browser (Karolchik et al., 2004). Salmon was launched with the following parameters : --numBootstraps 60 --libType A -- validateMappings.

For the second RNA-seq performed in NIR, IR and IR+TNFa conditions, we used nf-core/rnaseq (version 3.3) pipeline for RNAseq analysis (doi: 10.5281/zenodo.1400710), with the folowing additional parameters : --genome mm10 --clip_r2 14, and performed on the Core Cluster of the Institut Français de Bioinformatique (IFB) (ANR-11-INBS-0013).

#### Differential gene expression analysis

Statistical analysis was performed using R v3.5.1. Transcript expression levels were aggregated in Gene expression levels using tximport Bioconductor package (Soneson et al., 2015) v1.13.16. Deseq2 (Love et al., 2014) v1.22.2 method was used to identify differentially expressed genes between groups with a p-value threshold of 0.05.

#### Permutation test

To create the list of genes hosting an L1Md, BED files containing L1Md genomic localizations (reconstructed Repbase from (Walter et al., 2016)) were intersected with the refseq_curated database from UCSC. Permutation test (n=10,000) between lists of genes hosting an L1Md and DEG in IR *vs* NIR, or the same number of random genes (randomly extracted from Refseq without DEG) was performed using R studio and considered significant if *p<0.01*.

**Motif Enrichment Analysis** was performed using BaMM! web interface (Kiesel et al., 2018; Siebert and Söding, 2016) and *de novo* motif discovery module. Query motif was matched to known motifs using the hocomoco mouse database

**GSEA Analysis** was performed using Hallmark Gene Sets V7. To plot graphs, -log10 pValue is set to 4 when p<0.0001.

### ChIP-seq

#### Alignment

Human sequences were found in Mouse ChIP-seq reads. The contamination was removed with Xenome (Conway et al., 2012) v1.0.0. After contamination removal, ChIP-seq sequence reads were mapped to the Mouse genome build mm10 by using Burrows-Wheeler Aligner MEM algorithm (Li and Durbin, 2009) (BWA v0.7.17). The read group ID was attached to every read in the resulting alignment file (bam file) with the -R parameter, and shorter split hits were marked as secondary with -M. Samtools (Li et al., 2009) fixmate v1.9 was used to check mate-pair information between mates and fixed if needed on a name sorted bam file. The duplicate reads were tagged by samtools markduplicates using a position sorted bam file. Secondary alignments and unmapped reads have been filtered out and only properly paired reads have been kept. Two types of downstream analysis have been performed, with multimapped reads (mapping quality score >= 0) and one with uniquely mapped reads (mapping quality score >= 1). Cross-correlation scores (NSC and RSC) have been calculated by phantompeakqualtools package (Kharchenko et al., 2008; Landt et al., 2012) v1.2. DeepTools (Ramírez et al., 2016) bamCoverage v3.3.0 has been used to generate normalized bigwig files with the following parameters : --binSize 1 --normalizeUsing BPM -- extendReads –ignoreDuplicates. Then deepTools bigwigCompare was used to substract input signal from chip signal.

#### Peak calling

Areas in the genome enriched with aligned reads (also called peaks) were identified with MACS2 (Zhang et al., 2008) callpeak v2.1.2 with the following parameters : -f BAMPE -g mm10 -q 0.05 --broad --broad-cutoff 0.05 for H3K9me3 broad mark.

#### IDR (Irreproducible Discovery Rate) analysis

To measure the reproducibility between replicate experiments, we used the IDR method (Li et al., 2011) v2.0.4.2 with the following parameters : --rank q.value --random-seed 12345 --plot. Peaks with a global IDR score < 0.05 were selected and used for downstream analysis.

#### Peak annotation

Annotatr 1.8.0 (R3.5.1) was used for peak annotation.

#### H3K9me3 quantification and differential binding

To quantify H3K9me3 concentration at TE or promoters (−2kb;+1kb TSS), the Bioconductor package Diffbind (Ross-Innes et al., 2012) v2.10 was used in R v3.5.1. Paired-end mode was activated for read counting step with SummarizeOverlaps method. The default mapping quality threshold (mapQCth) was modified in 0 for multimapping analysis or 1 for unique mapping analysis. DBA_DESEQ2_BLOCK method was used to consider unwanted variable during normalization. Normalized H3K9me3 concentration at all TE loci from a same family/subfamily was summed to get a total H3K9me3 concentration per TE family. The age of a TE was calculated as in (Sookdeo et al., 2013): divergences were converted to time assuming a neutral rodent genomic substitution rate of 1.1%/MY.

Differential binding at peaks was identified with a p-value threshold of 0.05.

#### Heatmaps

To plot heatmaps of H3K9me3 enrichment at peaks, deeptools package v3.2.0 was used in R v3.5.1. The peaks (IDR<0.05) files obtained for NIR and IR conditions were first fused using bedops. A matrix was then built using ComputeMatrix tool in the scale-regions mode between the generated fused bed file and the corresponding normalized bigwig files after input substraction. A body length of 2.5kb (mean size of the peaks) was selected, as well as a 4kb distance upstream and downstream of the start and the end of the peak. We asked for a “–outFileSortedRegions” that gives the sorted bed file used for the heatmap. This sorted bed file was then used for genome coverage analysis, *i.e*.: identification of the presence of a given TE in each row after computing a matrix with the TE genome coverage bigwig.

#### TE genome coverage

To generate TE genome coverage, bedtools package v2.27.1 was used. –bga option on the genomeCoverageBed tool was used. The bedGraph generated were then converted to bigwig files using the bedGraphToBigWig tool.

#### Accession numbers

ChIP-seq and RNA-seq raw data deposition on Annotare (ArrayExpress) is ongoing.

### Cut&Tag

CUT&Tag-IT assay kit (Active Motif) was used on 3,000 HSC according to manufacturer’s instructions. Cells were incubated O/N with 0.25µg of H3K9me3 (C15410193-Diagenode).

Cut&Tag was analyzed as described as in https://yezhengstat.github.io/CUTTag_tutorial/index.html with the folowing parameters: Quality Control using FastQC(0.11.9) and MutiQC (1.10.1) ; - Bowtie2 (2.4.1) alignment to mm10 (UCSC genome):with the folowing parameters: --end-to-end --very-sensitive --no-mixed --no-discordant --phred33 -I 10 -X 700 ; - Remove duplicate using Picar (2.26.9) with the folowing parameters: --REMOVE_DUPLICATES true --VALIDATION_STRINGENCY LENIENT ; - Aligned read quality score set to 0 to keep all reads by using samtools (1.13) with the folowing parameters: -q 0 ; - Aligned reads were sorted and indexed using samtools (1.13) ; - Generating a coverage track (bigWig) using deeptools (3.5.0) with the folowing parameters: -bs 5 --normalizeUsing BPM ; - peack calling using macs2 (2.2.7.1) with the folowing parameters: -B --broad --broad-cutoff 0.1 -f BAMPE -g mm --max-gap 2000 --min-length 200 ; - profile plot for scores over genomic regions (mm10.rmsk.mod.L1Md.bed) using deeptools (3.5.0) with the folowing parameters: --beforeRegionStartLength 1000 -- regionBodyLength 5000 --afterRegionStartLength 1000 \ ; - Statistics of the CUT and TAG signal (R package Rseb 0.2.0) : Using as input a score matrix computed by deeptools’s computeMatrix, we Ploted the mean density profile of all condition with the SEM (standard error mean).

## Acknowledgments

We thank animal facility, the Genomic and the Imaging and Cytometry Platforms of Gustave Roussy for RNA sequencing and cell sorting and confocal analysis, respectively; S. Gregoricchio and Drs. C. Lobry and C. Guillouf (INSERM U1170, Gustave Roussy) for their advices for ChIP-seq analysis, D. Dubray for Statistical analysis, and Drs M. Goodhardt and D. Garrick (INSERM UMRS-1126, Paris) for helpful discussions. We also thank A. Teissandier and D. Bourc’his for providing us the reconstructed repeatMasker database. This work was supported by INSERM and grants from Ligue National Contre le Cancer (LNCC, Equipe labellisée EL2020) and Institut National du Cancer (PLBIO N°2020-095) to F. P., ARC Foundation (N° 20161204988), GEGLUC Paris-Ile de France (2017) LNNC (2018- 2019) and Agence National de la Recherche (ANRJCJC20-CE14-0018-01) to E.E.M. Y.P. and D.H. are recipients of fellowship from the Ministère de l’Enseignement Supérieur de la Recherche et de l’Innovation. A.S. is recipient of a fellowship from LNCC. F.H. is supported by INCA (PLBIO N°2020-095 to F.P.).

The authors declare no conflict of interest.

## Author contributions

Y.P., and E.E.M. performed the RNA-seq, ChIP-seq, ChIP-qPCR, experiments and analyzed the results. M.K.D., A.M.C and E.E.M. performed bioinformatic analyses. D.H., F.H., and A.S performed IF, qPCR and reconstitution experiments and analyzed the results. E.E.M. and F.P. designed and supervised the study, analyzed the results and wrote the manuscript

