## Supplementary material for "The NF-κB pathway controls H3K9me3 levels at intronic LINE-1 elements and hematopoietic stem cell gene expression in cis": Figures S1 to S5

### Figure S1

Pelinski et al.

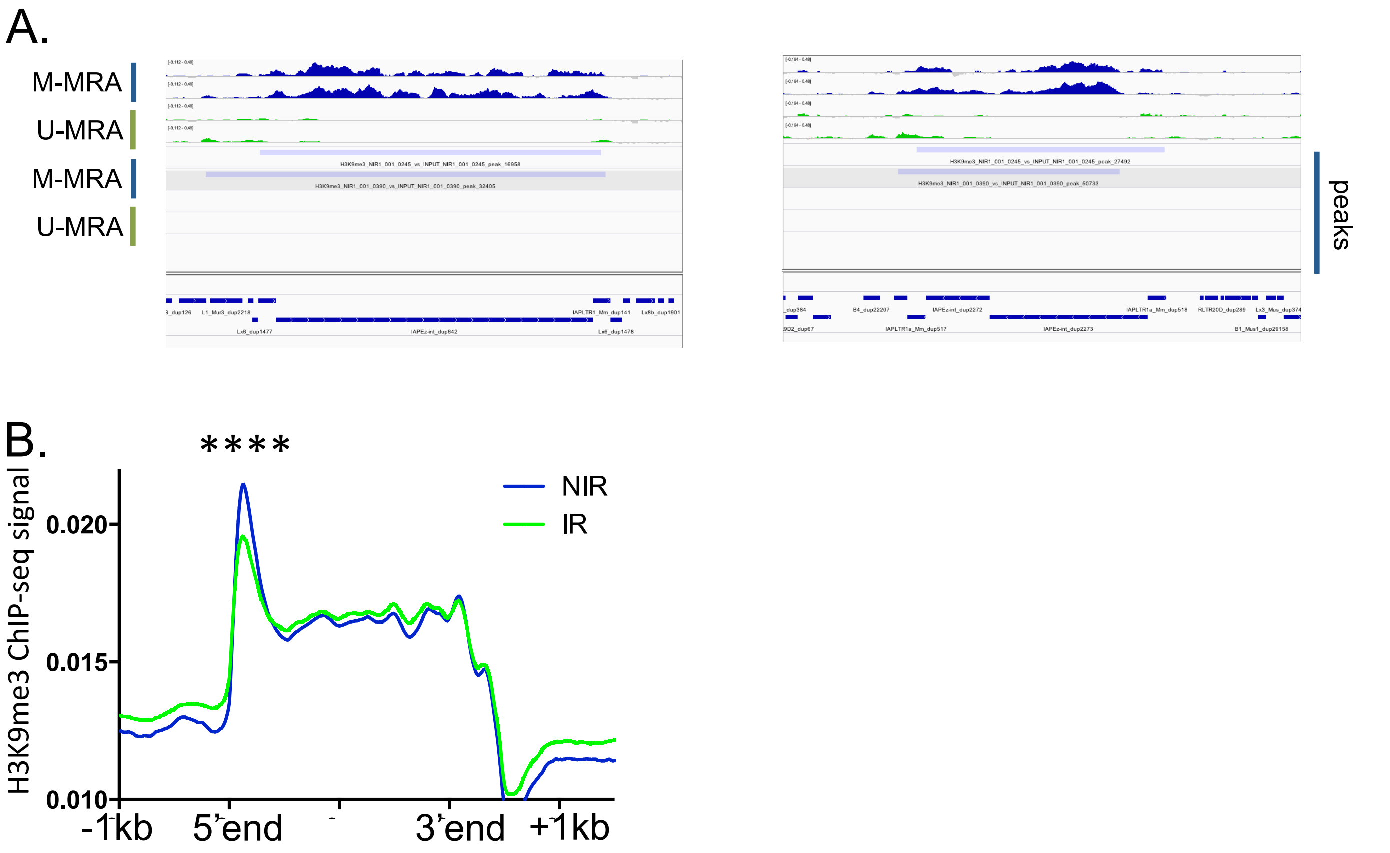


**Figure S1, related to figures 1 and 2. (A)** Integrative genomic viewer (IGV) visualization of H3K9me3 enrichment and peaks at 2 described loci with M-MRA or U-MRA analysis, as indicated. (left) chr2 :39209585-39320316 ; (right) chr6 :5271421-5288640 (Bulut-Karslioglu et al. 2014) **(C)** Plot profile representing H3K9me3 enrichment along the long L1Md sequences +/- 1kb flanking regions in NIR (blue) *vs* IR (green) conditions. ****p<0.0001 wilcoxon test

### Figure S2

Pelinski et al.

A.

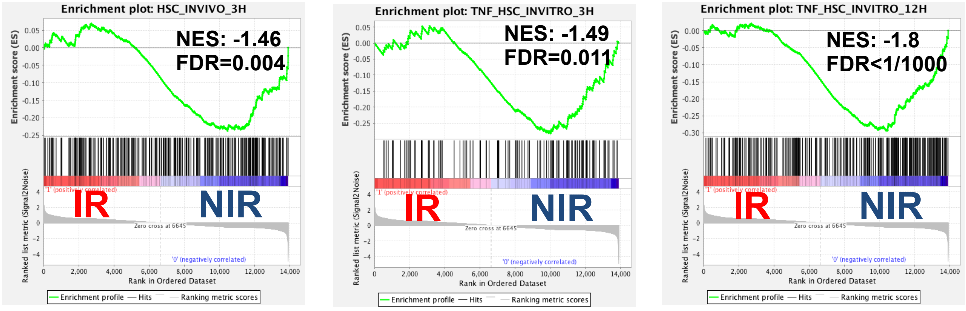


B.

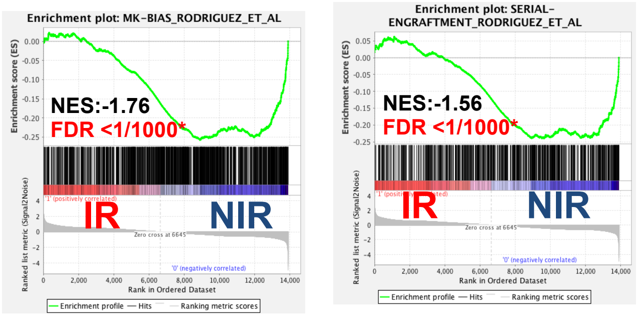


C.

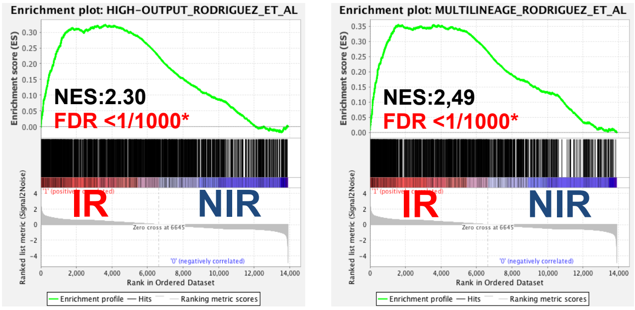

### Figure S2 continued

Pelinski et al.

D.

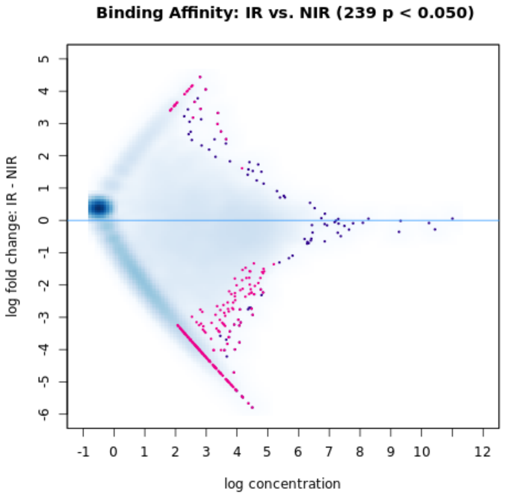


E.

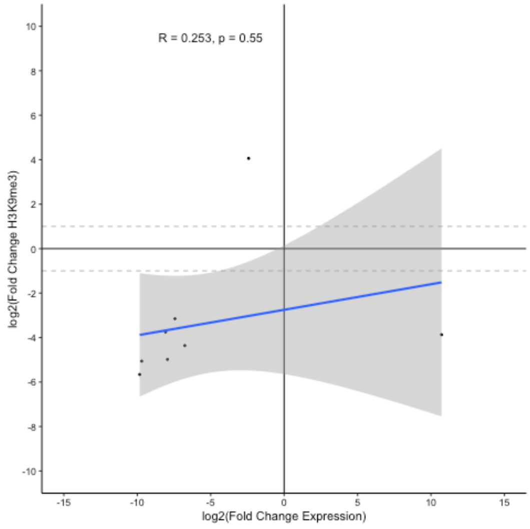


F.

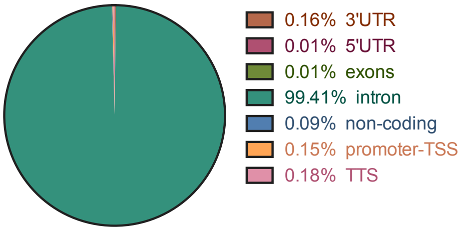


G.

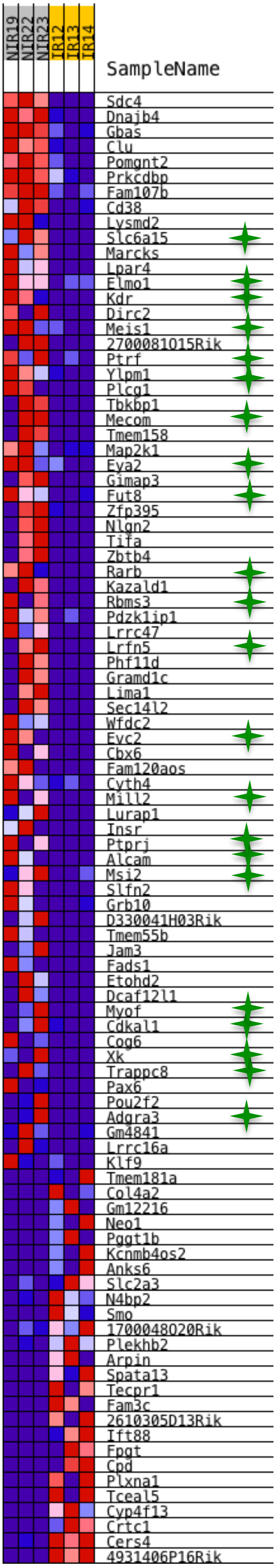


H.

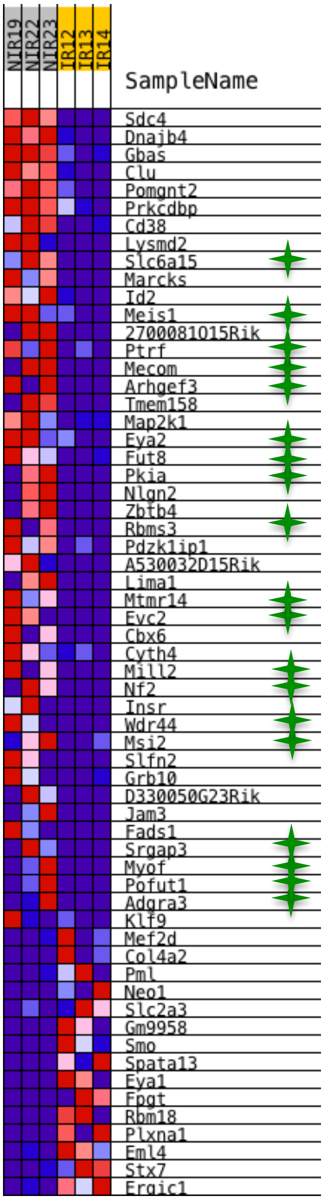

**Figure S2, related to figures 3, 4.** Enrichment plots in IR vs NIR conditions obtained from GSEA for **(A)** TNF signatures induced in HSCs after TNF-α treatment 3h *in vivo* and 3h and 12h *in vitro*; **(B)** MK-biased and serial engraftment HSC signatures; **(C)** High output and multilineage differentiation signatures

**(D)** Quantitative analysis of H3K9me3 enrichment at promoters (-2kb;+1kb TSS) performed by U-MRA. Non-significant (blue dots) and significant (*p<0.05*- pink dots) differential H3K9me3 enrichment at genes promoters are shown. **(E)** Correlation plot between H3K9me3 concentration at gene promoters *vs* gene expression at genes presenting both significant deregulation and differential H3K9me3 enrichment at their promoters upon IR (*p<0.05*). **(F)** Repartition of intragenic L1Md localization in the genes. TSS, transcriptional start site ; TTS transcriptional termination site. **(G,H)** Heatmaps of the expression of genes from the low-output **(G)** and MK-biased **(H)** HSC signatures that are significantly either upregulated (red) or downregulated (blue) in IR vs NIR. Green stars indicate the presence of an intronic L1Md in the downregulated genes.

### Figure S3

Pelinski et al.

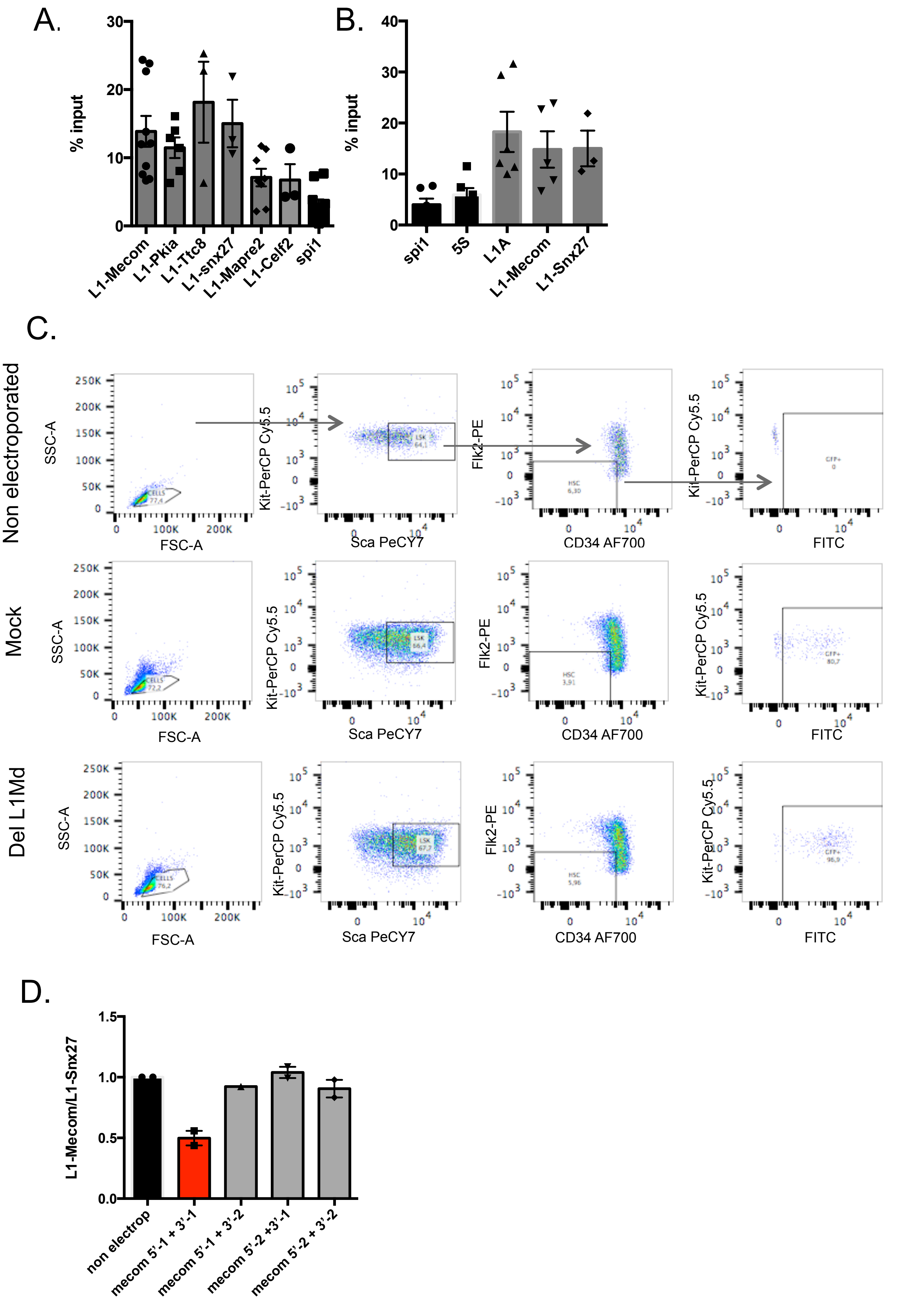

### Figure S3 continued

Pelinski et al.

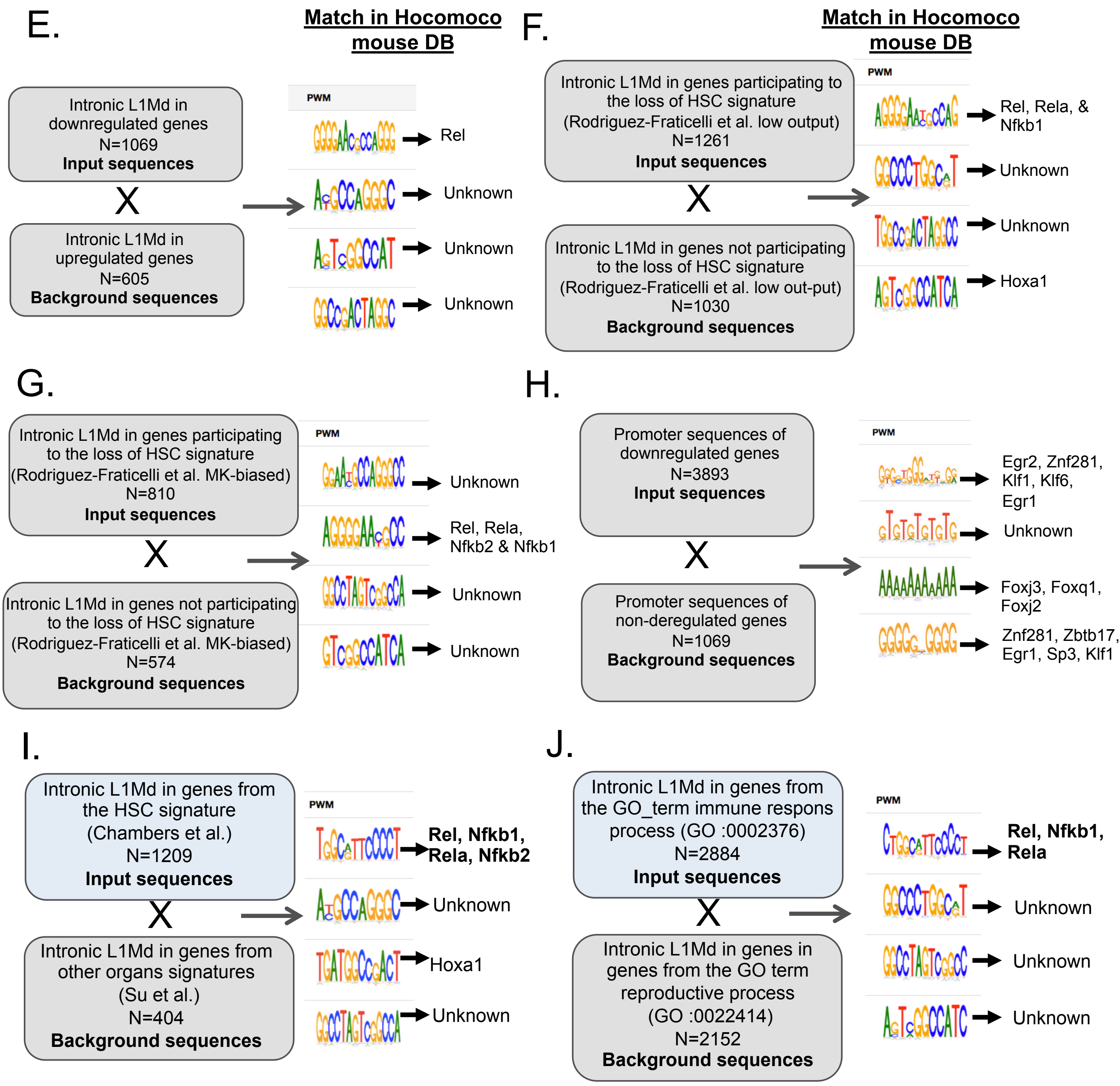

**Figure S3, related to figure 5**. H3K9me3 enrichment in the NIR conditions evaluated by ChIP-qPCR at long (>5kb) intronic L1Md promoters of the indicated genes compared to *Spi1* **(A, B)**, and at L1_A promoter, compared to repetitive 5S ribosomal RNA **(B)**. Results are expressed as the means +/- SEM of the percentage of input. Each dot represents a pool of 3 mice from 2-4 independent experiments. **(C )** gating strategy for electroporated siglo+ HSCs in mock or del L1Md conditions using non electroporated cells as a control for designing FITC gate. **(D)** To test the efficiency of the different combinations of gRNAs**,** DNA amplification of *Mecom* L1Md was assessed and normalized to the amplification of *Snx27* L1Md by RT-qPCR. In red the couple of guides that was selected for further analysis. **(E-J)** *De novo* motif discovery analysis performed with the BaMMmotif tool on L1Md sequences located in: **(E)** introns of downregulated genes *vs* updederegulated genes; **(F, G)** genes participating *vs* not participating to the loss of the low-output  or MK-biased HSC signatures; **(H)** promoter sequences (-2kb; +1kb TSS) from downregulated vs non deregulated genes; **(I)** L1Md located in genes from the HSC signature vs genes from other organs signatures (Kidney + liver + pancreas + testis + salivary gland + placenta) from Su et al. **(J.)** L1Md located in genes from the GO term immune response process (GO:0002376) vs genes from the GO term reproductctive process (GO: 0022414) . Enriched motifs were matched to known motifs using the Hocomoco mouse database.

### Figure S4

Pelinski et al.

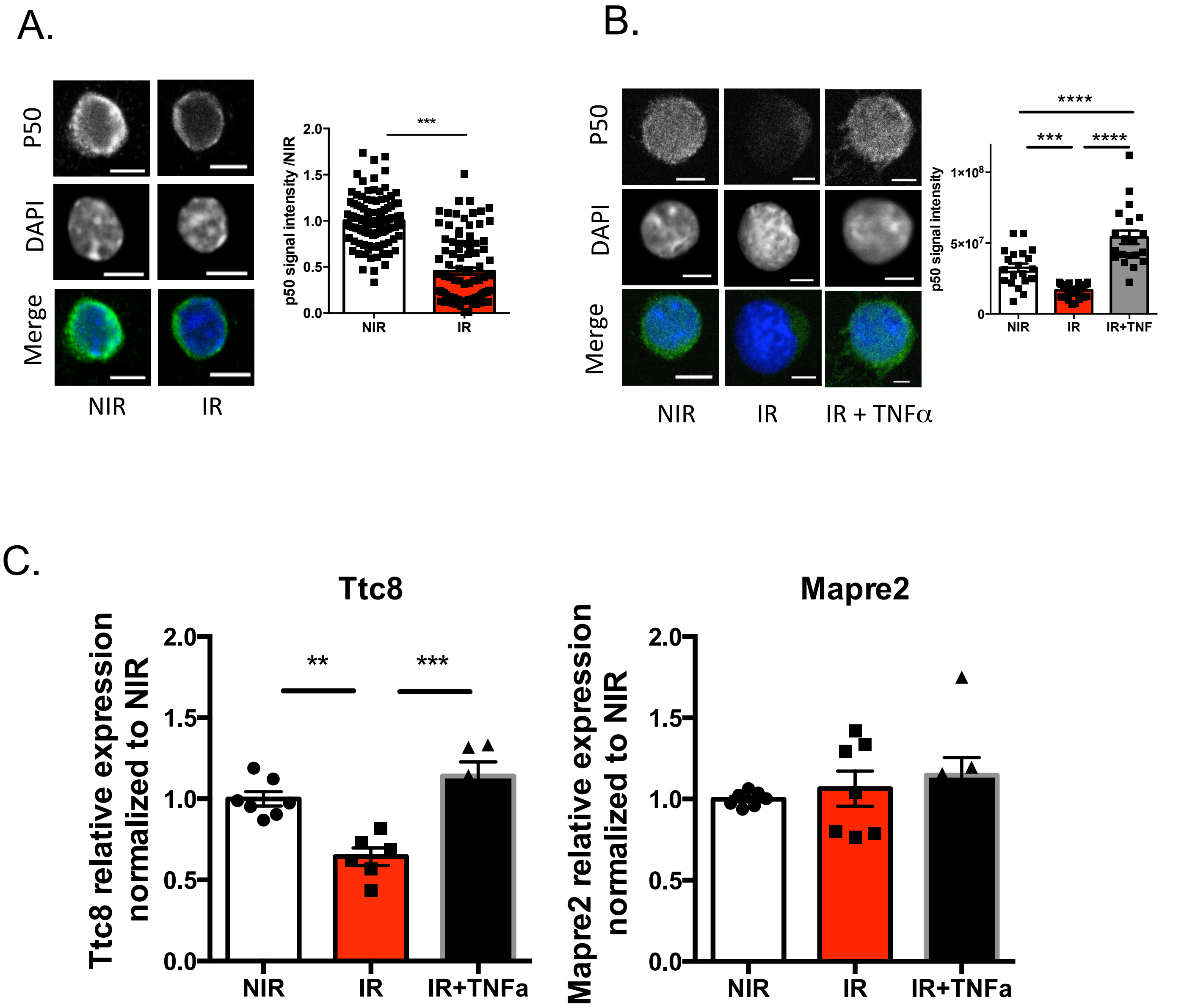


**Figure S4, related to Figure 6**. **(A, B)** Representative images and quantification of Nfkb1 protein mean IF intensity using the monoclonal anti-NFKB1 (p50) antibody clone from Santa Cruz Biotechnology. Bars, 5$\mu$M. Each dot represents a cell. Results are expressed as fold change from the mean value of the NIR condition from 2 independent experiments and represented as means +/- SEM.*** p<0.001 t-test for **(A)** or One-way ANOVA with Tukey's multiple comparison test for **(B)**. **(C)** *Ttc8* and *Mapre2* mRNA expression evaluated by RT-qPCR. Ct values were normalized to *Rpl32* and *Hprt*. Results are expressed as fold change from the mean value of the NIR condition and represented as means +/- SEM from 2 independent experiments. **p<0.01; *** p<0.001; one-way ANOVA Tukey's multiple comparison test.

### Figure S5

Pelinski et al.

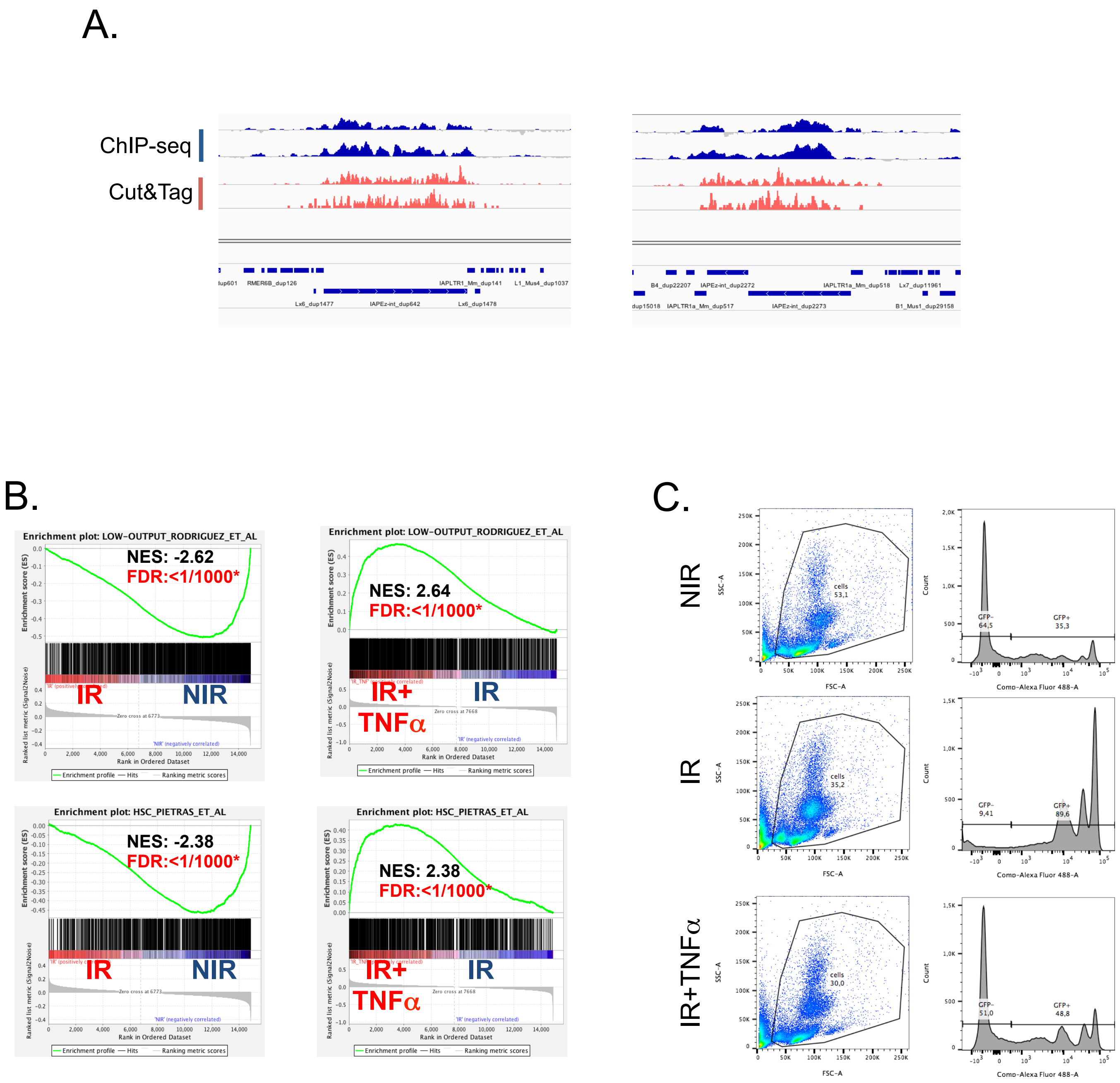

### Figure S5 continued

D.

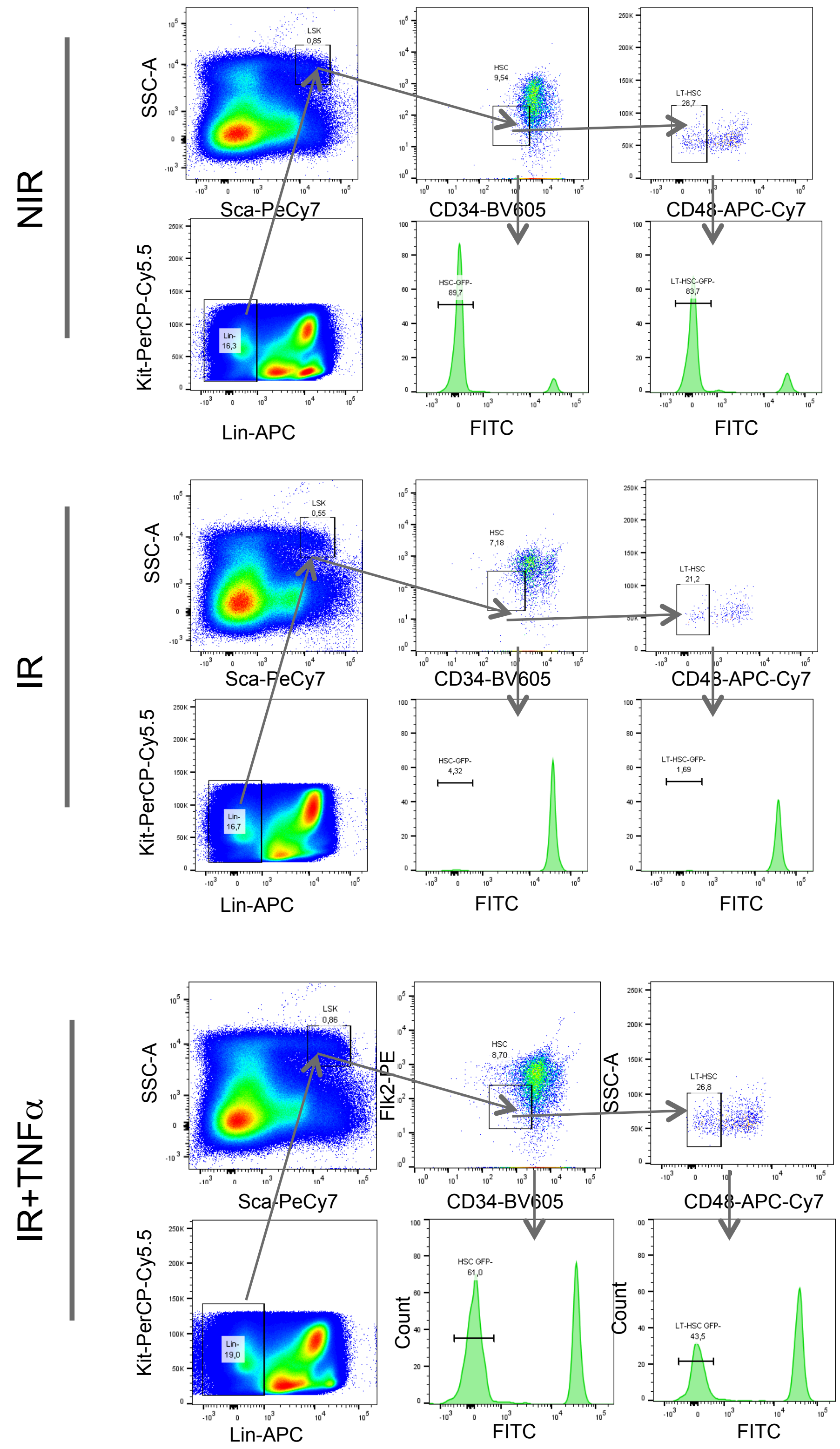


**Figure S5, related to figure 7. (A)** Integrative genomic viewer (IGV) visualization of H3K9me3 enrichment obtained by ChIP-seq and Cut&Tag M-MRA analysis at 2 described loci as indicated. (left) chr2 :39209585-39320316 ; (right) chr6 :5271421-5288640 (Bulut-Karslioglu et al. 2014). **(B)** Enrichment plots in IR vs NIR and in IR+TNFα *vs* IR conditions obtained from GSEA for low-output and LT-HSC signatures. **(C,D)** gating strategy to evaluate donor-derived GFP- vs recipient GFP+ cells in peripheral blood **(C)** and bone marrow HSCs **(D).**
